# Sensory constraints on volitional modulation of the motor cortex

**DOI:** 10.1101/2023.01.22.525098

**Authors:** Carmen F. Fisac, Steven M. Chase

## Abstract

Voluntary movement is driven by the primary motor cortex (M1), and individuals can learn to modulate even single neurons at will. Yet M1 also receives pronounced sensory inputs and contributes to sensory-driven motor responses. To what extent do these non-volitional signals restrict voluntary modulation of M1? Using a task in which the firing rate of a single neuron directly determines the position of a computer cursor along a visual axis, we assessed the ability of monkeys to modulate individual neurons under different sensory contexts. We found that sensory context persistently affected volitional control of single neurons in M1. For instance, visually rotating the biofeedback axis could render the same neural task effortless or problematic. Notably, extended training within or across days did not resolve this disparity. Our findings suggest that sensory context can limit the degree to which M1 activity is under volitional control.

## INTRODUCTION

The primary motor cortex (M1) is heavily involved in the generation of intentional movement^1^. Brain-computer interface (BCI) paradigms facilitate the study of sensorimotor control in a tractable environment^2^, making them powerful tools to probe the extent to which neurons in M1 can be volitionally modulated. Early studies showed that individuals can learn to reliably modulate even single neurons in M1 at will, if provided direct feedback of firing rate^3,4^. Further examination has shown that M1 activity can be flexibly dissociated from movement across a range of behaviors^5^. For instance, the strength of the correlation between M1 modulation and the bursts of muscle activity that ordinarily accompany it can be flexibly adjusted by conditioning on electromyography feedback, making it possible to both magnify the neuron-to-muscle connection^6^ or decouple it entirely^7^. Indeed, during BCI control, animals often stop moving the native limb altogether, even when the decoder is trained on overt movements^8,9^. Moreover, when neural activity is used to drive electrical stimulation of paralyzed muscle tissue, monkeys can learn to modulate individual neurons to control muscles regardless of their original correspondence with limb movement^10^. Similarly, subjects can volitionally control small ensembles of neurons in the motor cortex irrespective of prior directional tuning or co-modulation patterns^11–14^. The above body of literature suggests that there are few constraints—if any—on the volitional modulation of M1.

However, non-volitional signals are also present in the motor cortex, and may be powerful enough to compete with voluntary modulation. Anatomical studies describe prominent somatosensory inputs arriving at M1^15–17^, which can trigger salient sensory-evoked responses in the motor cortex^18–21^. Sensory-evoked modulation of M1 is not limited to somatosensation: visual perturbations of a BCI cursor can drive fluctuations in motor cortical activity unrelated to corrective cursor movements^22^, and visual feedback can influence BCI control depending on whether it is congruent or incongruent with proprioception^23^. There is also evidence of M1 involvement in reflexive motor responses. The long-latency response (LLR) precedes the earliest voluntary motor command when reacting to feedback disturbances during movement, and can be triggered by postural^24,25^ or visual^26,27^ perturbations. Unlike faster reflexive responses, LLRs have been shown to engage M1, as well as other cortical circuits fundamental to voluntary movement^28–33^. Finally, Collinger and colleagues have found that the context surrounding a movement can affect how that movement is encoded in M1: in human BCI clinical trials, they observed that the neural encoding of grasp changed depending on whether that grasp was performed in open air or around an object, to the point where it compromised grasp control^34^.

Together, these sets of observations beg certain questions. To what degree is neural activity in M1 under volitional control? How is voluntary modulation of M1 dependent on sensory context? To address them, we leveraged BCIs to design a paradigm that dissociates volitional and sensory drivers of neural activity. We trained three macaque monkeys to perform experiments in which the modulation of a single neuron’s firing rate in M1 dictated the position of a computer cursor along a one-dimensional axis. While the neural requirements were kept identical throughout a session, we isolated and altered aspects of the task’s sensory feedback in order to assess their influence on the subject’s ability to succeed at the task. We tested two manipulations of sensory context through this framework: axis orientation and location in the workspace.

Our paradigm revealed that certain sensory features can impact the ability to volitionally modulate single-cell activity in M1. Specifically, we found that the orientation of the cursor axis substantially impacted the ability to voluntarily modulate a majority of the neurons tested across all animals. While subjects would be able to easily and quickly modulate a given neuron to drive the cursor along the axis at one angle, they would immediately exhibit difficulty modulating it when the cursor axis was rotated, despite the neural task remaining the same across conditions. Changing the location within the screen where the cursor movement took place also had an impact on control, albeit to a lesser extent than orientation. Crucially, providing the subjects with days of additional training did not rescue their difficulty in controlling the cursor under certain sensory contexts. Combined, our findings suggest that the extent to which M1 activity is under volitional control is limited by sensory context.

## RESULTS

To assess the impact of sensory context on the ability to voluntarily modulate neurons within M1, we trained three rhesus macaques on the Feedback Alteration Single-cell Task (FAST). This BCI paradigm translates the firing activity of an individual neuron—termed the *command neuron* (CN)—into the real-time position of a computer cursor along a one-dimensional (1D) movement axis, and allows for flexible manipulation of task feedback to test the subject’s ability to voluntarily modulate the CN across sensory conditions (Fig. 1a).

**Fig. 1:**
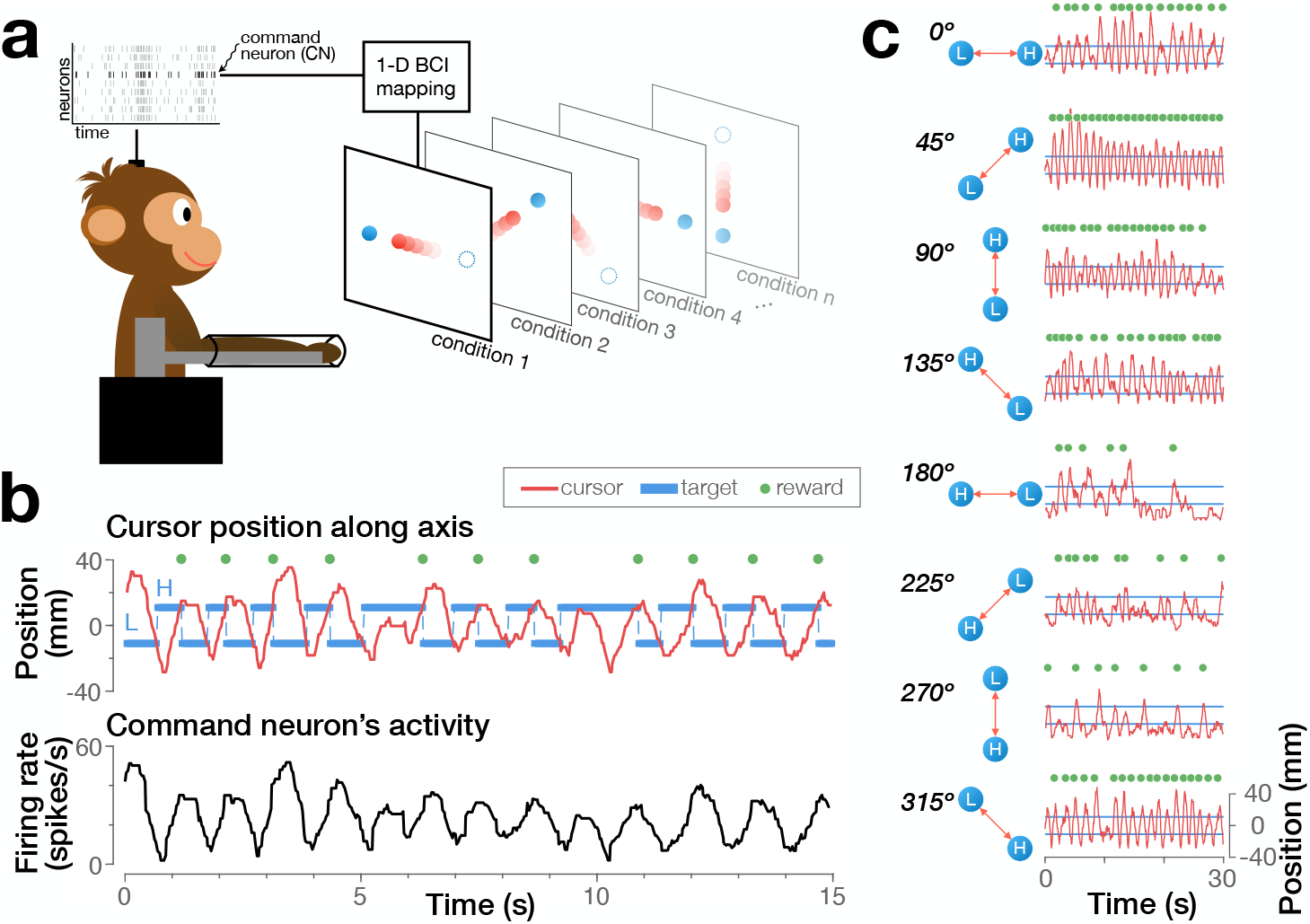
The FAST paradigm dissociates the neural requirements of the task from its sensory consequences. **a** A 1D BCI position decoder takes in the activity of a command neuron (CN) to move the cursor along an axis on the display. In the FAST orientation paradigm, each condition differs only in the orientation of that movement axis, while the neural requirements of the task remain constant throughout a session. A different neuron from the microelectrode array was chosen as each session’s CN. **b** The real-time position of the cursor (top) along the axis is determined by the CN’s spike count (bottom). In order to receive a reward, the subject must modulate the CN’s firing rate to drive the cursor from the low (L) to the high (H) target in under 2 s. **c** Example of the FAST orientation paradigm, in which the firing of the CN drives the cursor along axes that have different angular orientations in different blocks. Here we show the cursor position produced by the same CN during 30 s of representative performance for each condition. Note the variability in success rate across orientations. Data: session FOA06 (Monkey A).

In order to successfully complete a trial, subjects had to control the CN’s firing rate to drive the cursor along the axis and sequentially hit a low firing rate target and a high firing rate target on the screen within 2 s (Fig. 1b). Trials were presented continuously, with no inter-trial interval. Blocks of trials were characterized by the different sensory contexts provided; for instance, the orientation of the movement axis could vary across conditions (Fig. 1c). On a typical block, subjects were given 4 min to continuously modulate the CN and complete as many trials as possible on the same feedback condition. To calibrate each day’s FAST parameters, subjects first performed a short, standard 2D BCI center-out task to collect baseline firing statistics from the neural population (Methods).

### Characterizing volitional control

To determine whether sensory context restricts volitional modulation, we must first quantify volitional control. Good volitional control of an individual neuron is characterized by an ability to hit a wide range of firing rate values, and an ability to quickly transition from one firing rate to another.

To more fully probe the limits on controllability, the paradigm run on Monkeys N and R included an adaptive difficulty schedule that gradually increased task difficulty within individual blocks. Visually, the position of the high and low targets along the movement axis remained constant throughout a block; however, if recent task performance reached a certain threshold this would cause the gain between firing rate and cursor movement to shrink. This method expanded the dynamic range of neural modulation required to reach the visual targets on the screen (Fig. 2a), increasing task difficulty for future trials in the same block. Thus, multiple consecutive rewards in close succession resulted in a separation of the firing rate targets that the CN had to reach in order for the cursor to hit the same visual targets. The difficulty scaling was relative to the CN’s firing statistics and target rates were reset to their original values at the beginning of every new block (Methods).

**Fig. 2:**
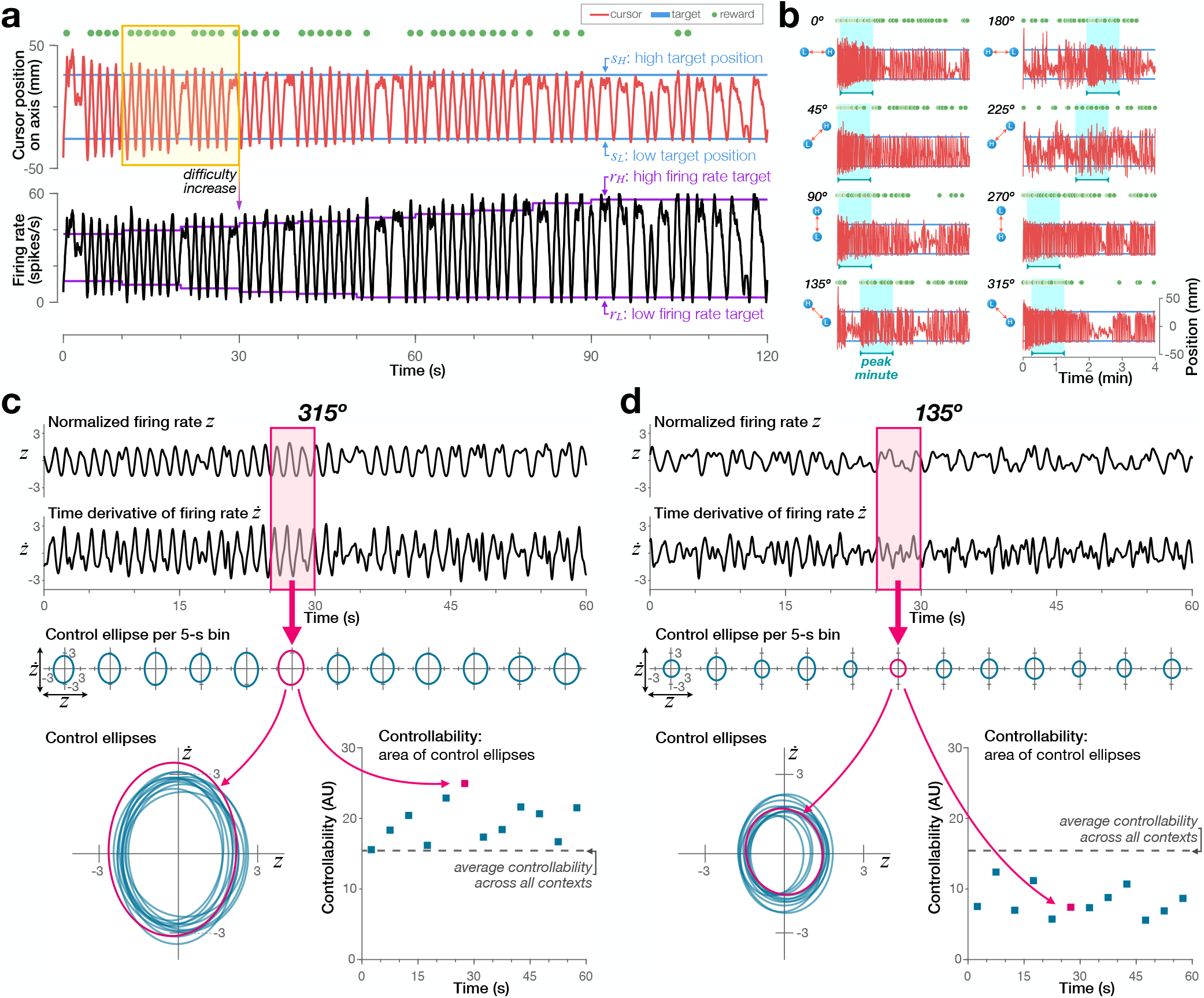
Controllability quantifies the ease with which a neuron’s firing rate can be modulated. **a** Cursor position and firing rate during the first 2 min of a block (Monkey N). Blue lines (top) show the position of the high and low targets along the axis; purple lines (bottom) show the firing rates required to hit those targets. We implemented an automated schedule that raised task difficulty by increasing the separation between the firing rate targets; the yellow box highlights one example in which recent performance triggered a difficulty increase. **b** A block’s peak minute (cyan shading) is defined as the 1-min, continuous interval of highest average success rate. Same conventions as **a** (Monkey N, same CN). **c** The CN’s firing rate and its derivative with respect to time are plotted against time during the peak minute of the 315° condition (top two rows). Controllability is computed in 5-s non-overlapping bins based on these values (pink box). Below each bin, we show its corresponding control ellipse: the 2-standard-deviation covariance ellipse between firing rate and its time derivative (teal). Controllability is defined as the area of these control ellipses (bottom). **d** Same as **c** for the 135° condition. Firing rates close to baseline or resistant to change indicate difficulty controlling the CN’s activity and produce low controllability values. At 315°, the subject was able to readily modulate this CN both widely and quickly, reflected respectively in the large fluctuations in firing rate and its derivative. At 135°, the animal was less adept at modulating the same CN. Data: session FON35 (Monkey N).

To quantify the volitional control of single cells in M1, we developed a *controllability* metric that captured both the magnitude and the speed with which the CN could be up- and down-modulated. We first identified the 1-min, continuous interval of best performance within each block based on success rate (Fig. 2b), and divided this *peak minute* into 12 non-overlapping, 5-second bins. Good control is characterized by large excursions in both firing rate and its derivative over this time period (Fig. 2c). In contrast, poor control is characterized by small deviations in these values (Fig. 2d). We captured these features with a *control ellipse*, formed by taking the area of a two-standard-deviation covariance ellipse between normalized firing rate and its time derivative (Methods). Larger areas subtended by the control ellipses, shown graphically as teal ovals in Fig. 2c-d, indicate better volitional control. Overall controllability for a particular sensory context is defined as the average of the control ellipse areas across the 12 bins within the peak minute.

### Sensory context limits volitional modulation of individual neurons in M1

We first tested whether the angular orientation of the movement axis impacts the ability to volitionally control a given neuron. To probe this, we varied the orientation of the movement axis throughout each session. If neurons are firmly tied to particular sensory contexts, then behavioral performance would change across directions. However, if voluntary modulation is truly dissociable from the effector, there should be little difference in controllability across orientation conditions.

We found that the orientation of the movement axis had a substantial impact on controllability. An example of this effect is displayed in Fig. 3, in which we show how the peak minute of each block maps to the CN’s controllability across all eight axis orientations. When the movement axis was angled at 135°, control was smooth and fluid, as marked by the sharp changes in firing rate and the wide dynamic range. This produced correspondingly large control ellipses for this orientation, leading to the high controllability values visible at the 135° condition in the central panel of the figure. In contrast, control was more coarse when the axis was oriented at 315°: in multiple instances, the subject had to make several attempts to reach the target after failing to drive the cursor far enough to receive a reward on the first try. These difficulties resulted in smaller control ellipses for the 315° condition, thereby producing the lower controllability values seen for this orientation in the central panel of the figure. By examining the differences in controllability across movement angles, we can weigh the impact that sensory context has on the subject’s ability to volitionally control the CN.

**Fig. 3:**
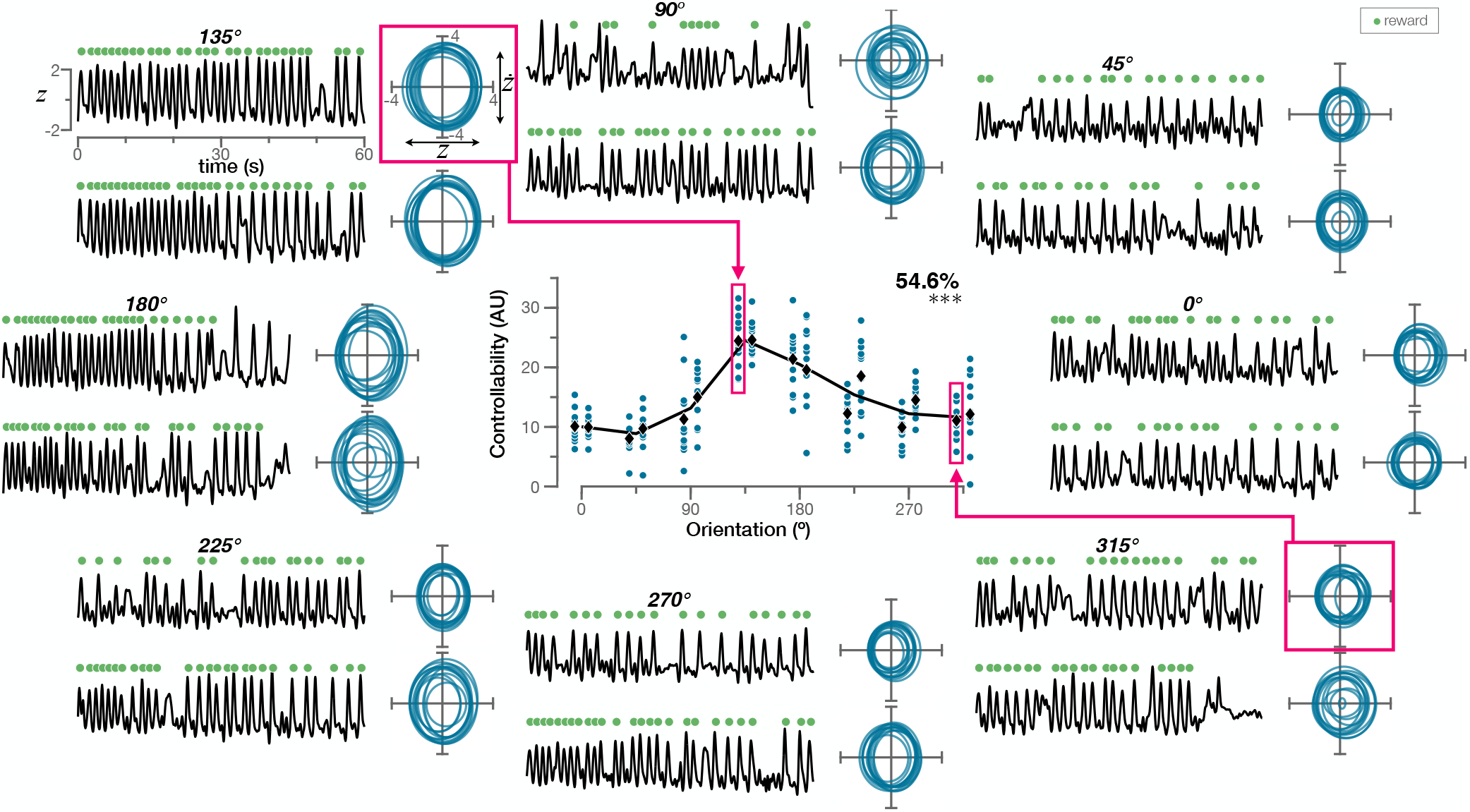
An example of the impact of orientation on single-neuron controllability. We assessed the ability to volitionally modulate the CN across different orientations by computing its controllability within the peak minute of performance on each block of trials. Block data are arranged on the periphery, based on sensory condition (each orientation was presented twice, non-consecutively). For each block, normalized firing rate during the peak minute is plotted against time (black), and its corresponding control ellipses are shown beside it (teal). Each ellipse represents the covariance between normalized firing rate (x-axis) and its time derivative (y-axis) during a specific 5-s bin within the peak minute. In the central panel, controllability is plotted against orientation. Controllability values from individual bins (teal) are grouped in the x-axis by condition, with the two blocks at the same orientation slightly offset horizontally from one another for visibility. Average controllability of a block is indicated by a black diamond; controllability averages per condition are connected by a line. We quantify the impact of orientation on the subject’s ability to volitionally modulate the CN by computing the fraction of the variance observed in controllability that is explained by orientation; this number (and a symbol indicating its statistical significance) is shown in the upper right corner of the central panel (one-way ANOVA; \*\*\**p* < 0.001). Data: session FON32 (Monkey N).

All three subjects displayed widespread changes in controllability as a function of movement orientation (examples from each subject shown in Fig. 4a, full set of CNs included in Supplementary Fig. 1). Indeed, a majority of CNs showed significant differences in controllability across conditions (one-way ANOVA, *p* < 0.05; N: 96%, 44 of 46 neurons; R: 100%, 16 of 16 neurons; A: 80%, 8 of 10 neurons).

**Fig. 4:**
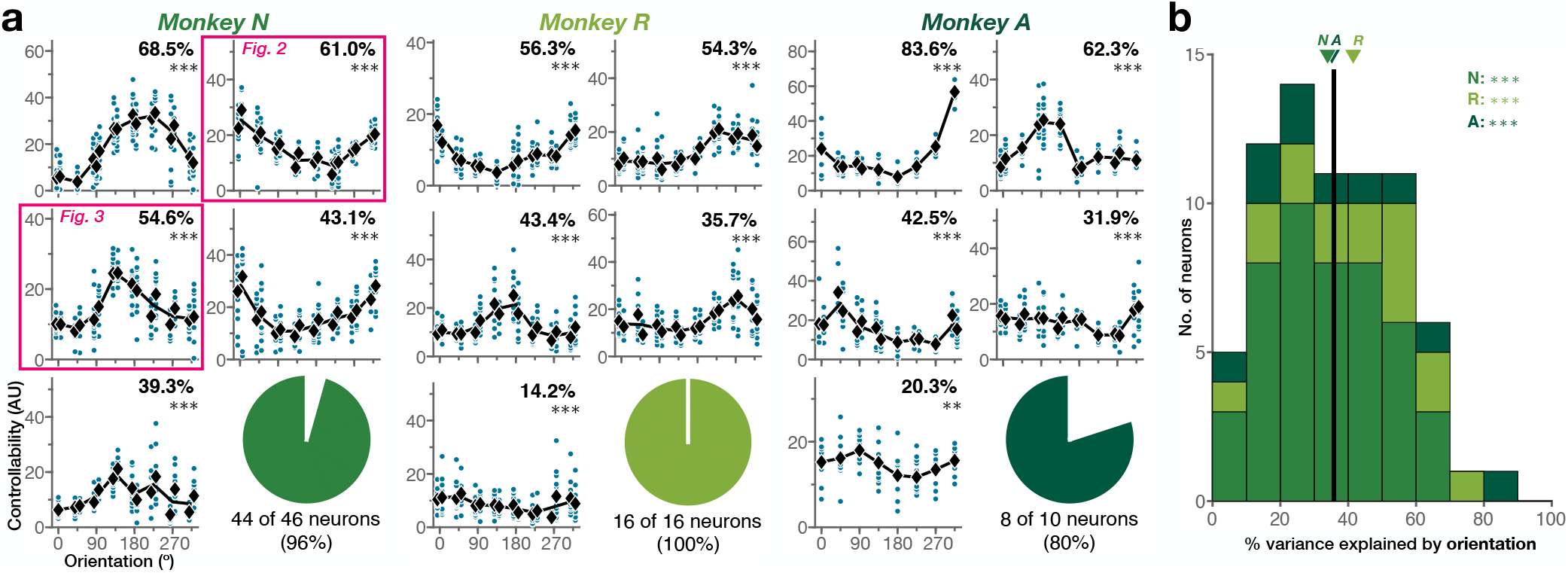
Volitional control is modulated by the orientation in which the 1D visual feedback is provided. **a** Controllability across orientations of example CNs for Monkey N (5 examples out of *n* = 46 neurons), Monkey R (5 examples out of *n* = 16 neurons) and Monkey A (5 examples out of *n* =10 neurons). For each neuron, individual controllability values (teal) and block averages (black) are shown; lines connect average controllability at each condition. Neurons are sorted by descending % variance explained by orientation, reported for each CN (one-way ANOVA; \*\*\**p* < 0.001, \*\**p* < 0.01). Pie charts indicate fraction of CNs that display significant controllability differences given orientation (one-way ANOVA; *p* < 0.05). **b** Distribution of the portion of variance in controllability explained by orientation across all neurons tested. Vertical line indicates average across subjects; arrows indicate subject averages. The portion of variance explained by orientation across CNs was statistically above chance for all subjects (Wilcoxon rank sum test; \*\*\**p* < 0.001).

We can quantify these changes in controllability by computing the variance in a CN’s controllability that is explained by orientation. Disparities in controllability across blocks were largely accounted for by the change in the angle of the movement axis (Fig. 4b), with an average of 35.8% of variance in controllability being explained by orientation across animals (N: 33.9%, R: 41.5%, A: 35.2%). For all three subjects, comparison with a null distribution in which controllability values were scrambled across orientations showed that the portion of variability explained by orientation was significantly greater than chance (Wilcoxon rank sum test; N: *T* = 3,158, *p* = 1.8 × 10^−15^; R: *T* = 389, *p* = 2.7 × 10^−6^; A: *T* = 152, *p* = 4.4 × 10^−4^).

Certain factors might impact neural modulation over time, such as a gradual decline in motivation throughout the session^35^. To control for effects of this timescale, we tested whether a linear change in a given CN’s controllability over time might explain the interaction between orientation and controllability that we had observed. We found that the dependence of controllability on orientation could not be explained by the passage of time across blocks (Supplementary Fig. 2).

We next evaluated whether volitional control was modulated by the overall location of the movement axis across the workspace (Fig. 5a, left). To do this, we trained two subjects on the FAST location paradigm (Monkey N, *n* = 20 neurons; Monkey R, *n* = 3 neurons), in which we fixed axis orientation at a particular value and instead varied the coordinates of the axis origin within the display (Fig. 5a, right).

**Fig. 5:**
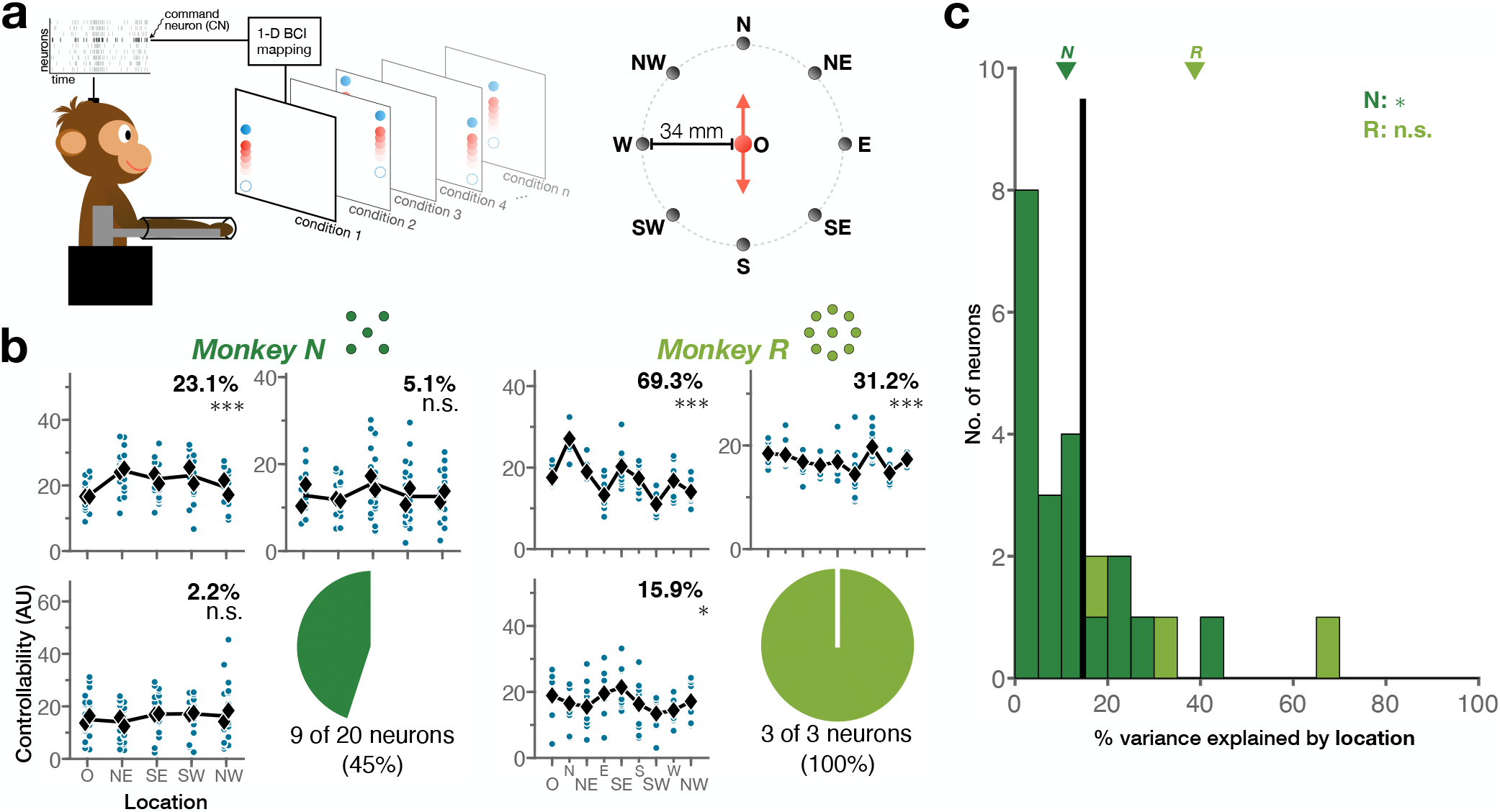
Location of the movement axis can also interact with volitional control. **a** In the FAST location paradigm, each condition is defined by the location of the movement axis within the workspace, tested for a single axis orientation at a time (left). Of the nine possible axis locations (right), Monkey N was tested on a subset of five locations (O, NE, SE, SW, NW); Monkey R was tested on all nine locations. **b** Controllability across locations of example CNs for Monkey N (3 examples out of *n* = 20 neurons) and Monkey R (*n* = 3 neurons), for one orientation. Individual controllability values (teal) and block averages (black) are shown for each neuron; lines connect average controllability at each condition. Neurons are sorted by descending % variance explained by location at this orientation, reported for each CN (one-way ANOVA; \*\*\**p* < 0.001, \**p* < 0.05, *n.s. p* ≥ 0.05). Pie charts indicate fraction of CNs that display significant controllability differences given location (one-way ANOVA; *p* < 0.05). **c** Distribution of the portion of variance in controllability explained by location, given one orientation. Vertical line indicates average across subjects; arrows indicate subject averages. The portion of variance explained by location across CNs was statistically above chance for Monkey N but not Monkey R (Wilcoxon rank sum test; \*\*\**p* < 0.001, *n.s. p* ≥ 0.05).

Both animals showed some differences in controllability due to axis location (examples from each subject shown in Fig. 5b, full set of CNs included in Supplementary Fig. 3). Yet, in contrast to orientation, a lower fraction of CNs displayed a significant impact due to location (one-way ANOVA, *p* < 0.05; N: 45%, 9 of 20 neurons; R: 100%, 3 of 3 neurons). Overall, location had a modest effect on controllability compared to orientation (Fig. 5b), with an average of 14.6% of variance in controllability being explained by the location of the movement axis across animals (N: 11.0%, R: 38.8%). The portion of variance in controllability that is explained by location was significantly greater than chance for Monkey N but not for Monkey R (Wilcoxon rank sum test; N: *T* = 500, p = 0.016; R: *T* = 15, *p* = 0.1), the latter likely limited by the lower sample size.

To control for any influence on the measured location effects due to axis orientation, for each CN we also tested controllability across locations on a second orientation. We found that the impact of location on controllability was comparable across both orientations (Supplementary Fig. 3), and again could not be explained by the passage of time across blocks (Supplementary Fig. 4).

### Further training does not eliminate the sensory dependence of volitional control

In the FAST paradigm, the neural requirements of the task are held constant while the sensory feedback varies. Of course, subjects may not be aware of this. Perhaps the struggle to control the CN in certain conditions was in part due to the animal not realizing that the neural task remained unchanged. We reasoned that, if this were the case, additional training at more troublesome conditions would result in improved control and a reduced interaction between sensory context and controllability.

To test this, we developed the FAST focused training paradigm, in which we followed a set of 4-min blocks across all orientation conditions with two *focus* blocks: two 15-min blocks at a single orientation. We then tested the subject’s controllability again across all orientation conditions (Fig. 6a). By comparing controllability curves before and after the 30 min of focused training at a single orientation, we can evaluate whether extended practice in an individual sensory context results in better volitional control of the neuron and less interaction between axis orientation and controllability.

**Fig. 6:**
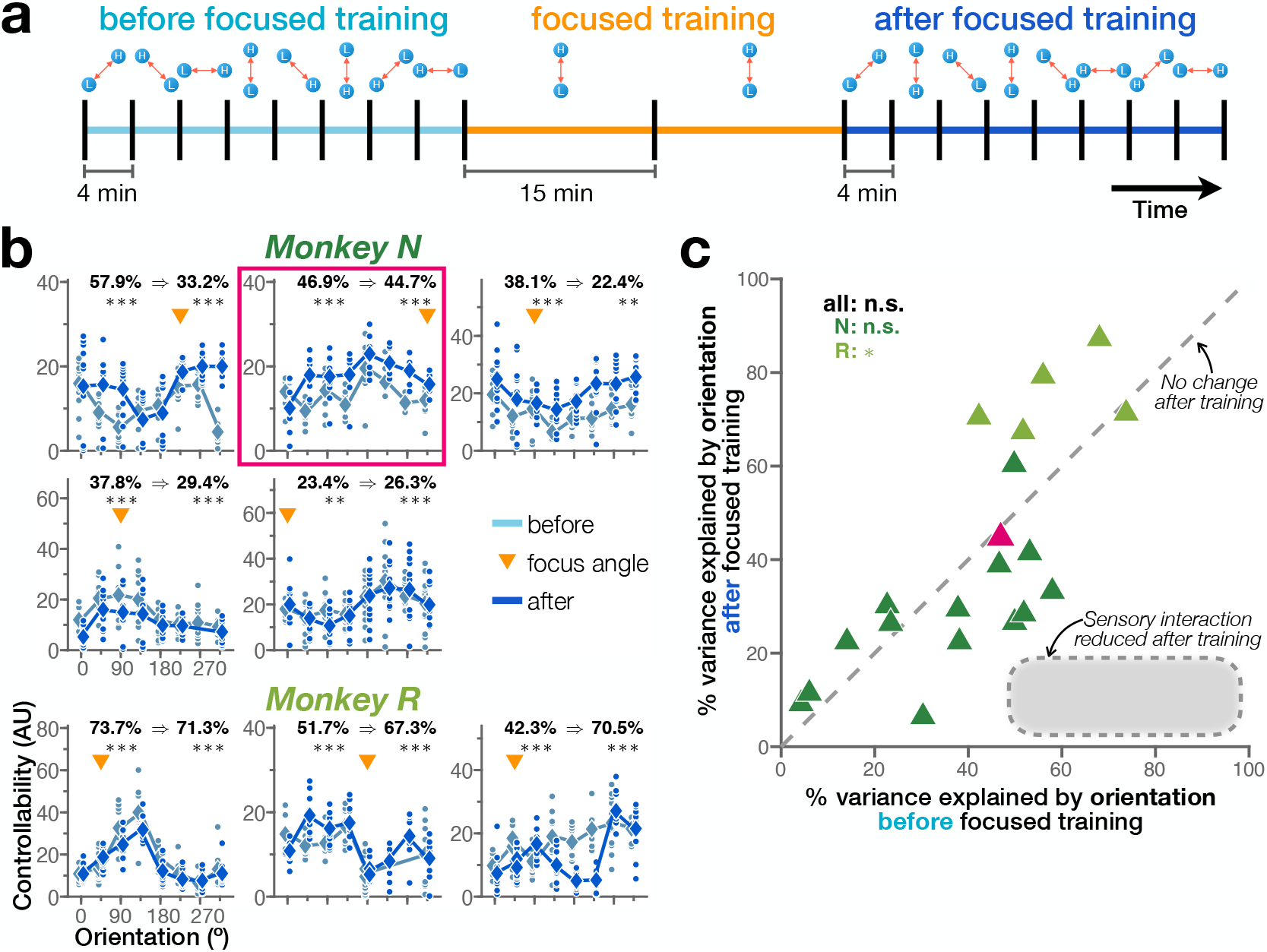
Focused training on a “difficult” sensory context does not extinguish orientation dependence. **a** The FAST focused training paradigm provides additional practice time in challenging conditions. An initial evaluation of controllability across all 8 conditions is followed by an extended period of training in a single orientation. Controllability across all sensory contexts is then assessed once again. **b** Controllability of example CNs across orientations before and after focused training for Monkey N (5 examples out of *n* =15 neurons) and Monkey R (3 examples out of *n* = 5 neurons). Controllability values (dots), block averages (diamonds) and lines connecting average controllability at each condition, before and after focus blocks, are shown for each CN. Variance explained by orientation is given as *before ⇒ after* (one-way ANOVA; \*\*\**p* < 0.001, \*\**p* < 0.01); neurons are sorted by descending variance explained before focused training. **c** Comparison of controllability variance explained by orientation, before and after focused training. Overall, there was no significant attenuation of the interaction between orientation and controllability (paired-sample t-test; \**p* < 0.05, *n.s. p* ≥ 0.05).

Prolonged practice in a condition that had proved challenging earlier in the session did not neutralize the interaction between orientation and controllability. Neither subject showed large differences in controllability before versus after the focus blocks (examples from each subject shown in Fig. 6b, full set of CNs included in Supplementary Fig. 5). One example of this is highlighted in Fig. 6b. This CN displayed below-average controllability at the 315° condition during the baseline block. After the subject was given two 15-min blocks at 315° to practice control in this sensory context, there was a visible improvement in controllability at this orientation. However, this effect was not restricted to the trained condition exclusively: prolonged training in the 315° orientation resulted in an overall increase in controllability across all sensory contexts. This sign suggests a learning effect that was independent of sensory context. Across subjects, we found no overall significant change in variance explained by orientation before versus after the extended practice given during the focus blocks (Fig. 6c); in fact, for Monkey R this additional training actually slightly *increased* the dependence of controllability on orientation.

To further evaluate the robustness of the orientation effect, we expanded our study of single-neuron controllability across multiple days with Monkey N. The FAST multi-day training paradigm provided the subject with 5 days of practice at the orientation task to control a single CN. By extending training in all orientations over several days, subjects were allowed multiple sessions to control the same neuron and refine their ability to modulate its activity across conditions.

We found that training for several days also did not eliminate the interaction between orientation and volitional modulation of individual neurons (one example shown in Fig. 7a, full set of CNs included in Supplementary Fig. 6). While we observed a slight weakening of the effect during the first 2 days of repeated training with the same CN (paired-sample t-test; *t* = 3.61, *p* = 0.009), additional training in the following days did not further attenuate the impact of orientation (paired-sample t-test; *t* = −0.09, *p* = 0.933), and the interaction between movement angle and controllability never disappeared (Fig. 7b). We did note an increase in the overall controllability of individual neurons with multi-day training when averaged across all orientations (Fig. 7c). These changes paralleled those in the attenuation of the interaction between orientation and controllability across days: average controllability increased between day 1 and day 3 (paired-sample t-test; *t* = −3.40, *p* = 0.011), but then reached a plateau (paired-sample t-test; *t* = −0.29, *p* = 0.782). Again, monotonic changes in controllability throughout each training session did not explain the interaction between orientation and controllability as a product of the passage of time across blocks (Supplementary Fig. 7).

**Fig. 7:**
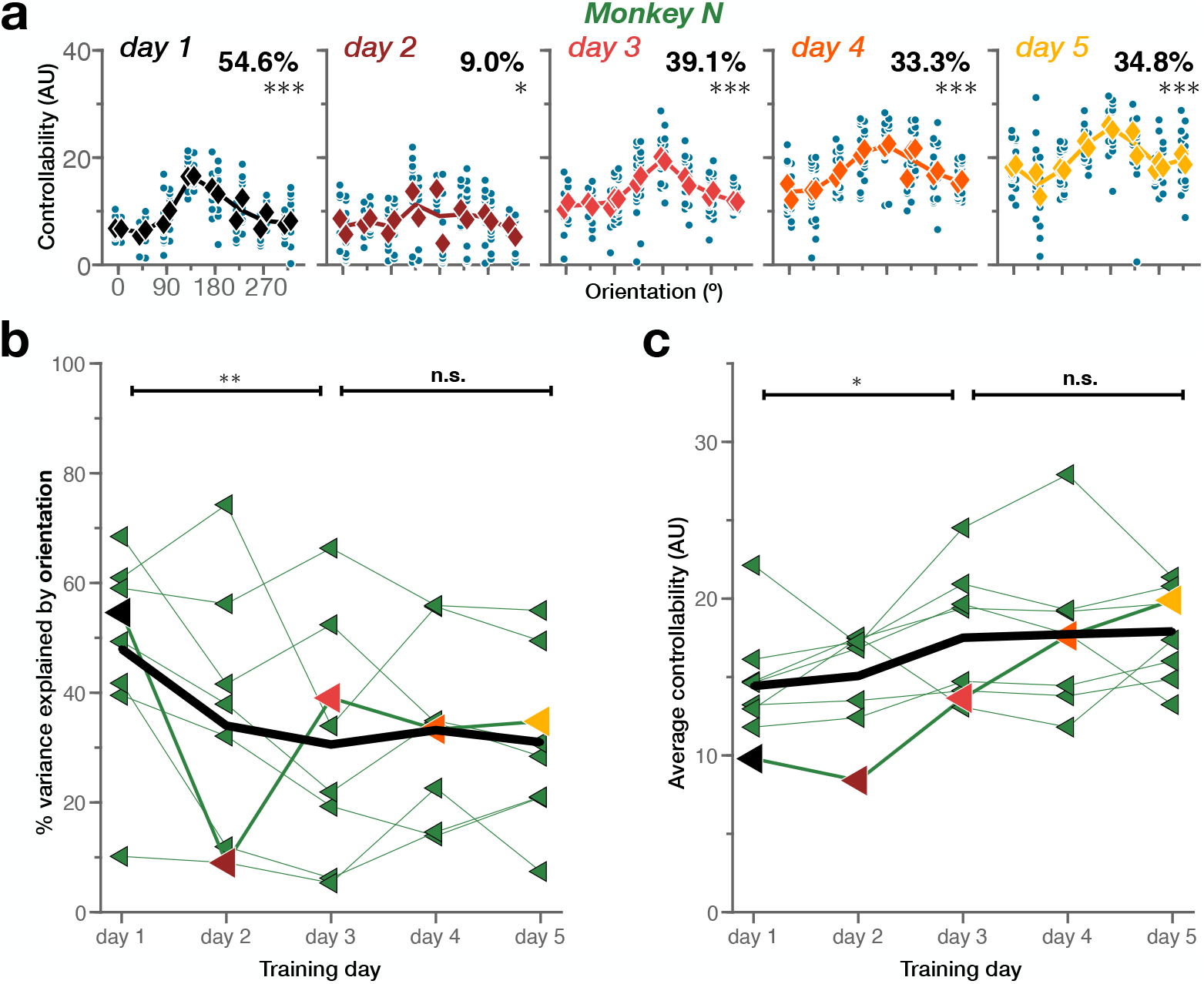
Training over several days is not sufficient to extinguish orientation dependence. **a** Controllability of one example CN across orientations per day for Monkey N (*n* = 8 neurons). Individual controllability values (teal) are shown; average controllability per block (diamond) and lines connecting average controllability at each condition are color-coded by training day. Variance explained by orientation is reported for each day (one-way ANOVA; \*\*\**p* < 0.001, \**p* < 0.05). **b** Portion of variance in controllability that is explained by orientation across days. Lines represent the different CNs (the example from **a** is highlighted); black trace indicates average across CNs. **c** Average controllability across days. Same conventions as **(b)**. Overall, early training (days 1–3) showed significant differences in both % variance explained and average controllability, while late training (days 3–5) did not (paired-sample t-test; \*\**p* < 0.01, \**p* < 0.05, *n.s. p* ≥ 0.05).

## DISCUSSION

We investigated whether the ability to volitionally modulate the activity of single neurons in M1 was affected by changes in sensory context, i.e., the manner by which feedback of firing rate was provided to the subject. We found that sensory context can indeed impose constraints on the degree to which individual neurons can be volitionally controlled. Across animals, controllability was heavily impacted by the orientation of cursor movement: subjects showed consistent differences in success when attempting to modulate individual neurons at various movement angles. Movement location within the workspace had an influence on single-neuron controllability as well; location effects were, however, less pronounced than those of orientation. Finally, we found that the interaction between sensory context and volitional controllability was robust to additional training. Providing animals with extended practice within the same session at an orientation that they had found problematic earlier was not sufficient to overcome these differences in controllability, and neither was granting a subject multiple days to improve control of a single neuron.

Why would movement location within the workspace have less of an impact on controllability than movement orientation? One potential explanation is that the magnitude of the displacement across location conditions was limited in our paradigm to accommodate all axis orientations and still guarantee that cursor movement remained within the confines of the display. Nevertheless, the lesser impact of movement location we observed is consistent with previous reports that cursor position in a BCI setting had only minor correlates in motor cortical activity and did not interfere with performance during neuroprosthetic control^36^.

In this study, we manipulated feedback exclusively within the bounds of a single sensory modality: vision. Of course, other forms of sensory feedback might also influence volitional modulation in the motor cortex. Postural changes, for instance, have been shown to sway directional tuning of individual neurons in M1^37,38^; the impact of posture could extend to controllability as well. Auditory signals have also been shown to evoke activity in the motor cortex. For example, human participant studies have found that speech sounds can elicit activity in motor areas^39,40^. Individual neurons in M1 have also been observed to change their firing as a function of object-related information during reach-to-grasp naturalistic movement^41^, to the point that whether a reach involves an object or not can lead to differences in M1 activity that disrupt BCI control of a robotic arm if not explicitly accounted for in decoder design^34^. Alongside these sensory effects, other variables not directly linked to sensory stimuli could potentially influence single-neuron controllability. Fluctuations in internal states—from attention to emotional condition—have been observed to modulate ongoing cognitive processes like decision-making, motor learning or sensory perception^42–44^, and can be detrimental for BCI decoding performance^35,45–47^. Further examination would be required to determine whether these factors might also limit our ability to volitionally modulate individual neurons in M1.

Even several days of additional training did not lead to a sizable decrease in the interaction between sensory context and controllability. This suggests that one of two possibilities is true: either this interaction is very hard to unlearn because of hard-wired constraints within the nervous system, or this interaction *can* be unlearned, but subjects did not stumble onto the proper solution even with days of practice. Both cases are interesting. If the first is true, and the interaction is hard to unlearn because of hard-wired constraints, it suggests we need to better understand how to measure these constraints. Previous research has identified key factors that govern whether a BCI mapping should be learned quickly or slowly^48,49^. Typically, a BCI mapping that only requires existing patterns of neural activity to be mapped to new movements are learned fast, showing substantial improvements on a single day of training. In contrast, BCI mappings that require new patterns of activity are learned slowly^50,51^. The FAST paradigm, by definition, only requires patterns of neural activity that the subject has already demonstrated in another context, and thus by prior theory should be learned quickly. This would imply we need new theory that can account for sensory interactions in the generation of motor neural activity patterns. If, on the other hand, the latter possibility is true, then there are no hard-wired constraints affecting volitional control across sensory contexts. This implies that subjects simply never came across the mental solution that would have enabled them to control the cursor in the new sensory context. If this is the case, it suggests that failures in learning could be a major factor limiting performance in the clinical implementation of BCI devices, and paradigms will need to be developed to guide subjects to the optimal neural patterns necessary to control a particular device.

While neither focused nor multi-day training were able to eliminate the interaction between sensory context and volitional control, the focused training paradigm did reveal some interesting features. It might have been the case that training at a single orientation would result in improvements in control along that orientation but not others. This is the case for visuomotor rotation learning, in which training to make movements to a single target under rotated visual feedback results in clear improvements to movements in that direction, with limited generalization to other target directions^52^. However, the most common response we observed was an increase in controllability that generalized across all directions, and thus maintained the interaction with sensory context. This is more akin to visuomotor gain learning^52^. These results imply that, whatever mechanism is responsible for the improvement in volitional control that comes with practice, it is not tied to the sensory context.

The notion that sensory context can impose bounds on our ability to voluntarily modulate M1 can have direct implications for the development of clinical neuroprosthetic devices. If the activity of individual neurons can be tied to particular sensory outcomes, these relationships could have an effect on how readily a BCI mapping can be learned. In other words, building decoders that are consistent with the sensory interactions in the population could constitute an additional factor in dictating how learnable a BCI mapping can be. Further study of this sensorimotor crossroads in M1 could enable decoders to take advantage of native neural pathways and dodge configurations that prove detrimental for BCI control.

The precise nature of volitional control is still very much an open question. Our results suggest that volitional modulation of M1 is subject to constraints imposed by sensory inputs. This would be consistent with a model of cortex in which sensory and volitional signals impinge on the same neurons, resulting in a mixing of sensory and volitional signals at the level of individual cells. If this holds true for other cortical areas, this framework would be consistent with the observation of volitional modulation across diverse areas of the brain, including primary sensory areas^13,53–60^, as well as explaining sensory-driven activity in motor areas. If both volitional and sensory signals coexist across the entire cortex, perhaps the key difference between sensory and motor areas is the degree to which they are under volitional control versus sensory-obligate modulation.

## METHODS

### Experimental procedures

All animal procedures were approved by the Institutional Animal Care and Use Committees of Carnegie Mellon University and the University of Pittsburgh.

#### Neural recordings

Three male rhesus macaques (*Macaca mulatta*) were each implanted with a 96-channel, 1.5-mm-shank recording microelectrode array (Blackrock Microsystems), inserted into the primary motor cortex (M1) using standard procedures^61^. Arrays were visually placed on the caudal convexity of the precentral gyrus at the level of the spur of the arcuate sulcus of the hemisphere contralateral to the trained arm, aiming for the proximal arm representation. Implant surgery occurred only after animals were trained to perform standard center-out reaching tasks in a virtual reality environment.

Microelectrode array voltage signals were amplified (1,000x – 32,000x), band-pass filtered (250 Hz – 8 kHz) and processed with a multichannel acquisition processor system (Plexon Inc., Monkeys N and A; Blackrock Microsystems, Monkey R). Waveform data was converted to spiking events by threshold crossing, and snippets that crossed the threshold were manually sorted online to isolate neurons for BCI decoding. During all recordings described in this study, the subject’s arms were lightly restrained. Subjects tended to exhibit minimal overt arm or hand movements while performing the tasks.

#### Task flow

The FAST paradigm dissociates the neural requirements of the task from the sensory context, and allows us to evaluate a subject’s ability to volitionally modulate the activity of individual neurons across a range of sensory feedback conditions.

Cursor position along a one-dimensional (1D), invisible axis was determined by the activity of a single command neuron (CN). Each trial required the subject to push the cursor to 2 targets appearing in succession on the movement axis: a *low* firing rate target, followed by a *high* firing rate target. Cursor and target radii were 4 mm each. No hold time was enforced at either target; subjects merely had to exhibit a firing rate that made the cursor pass through the target boundary for the hit to count, which would cause the target to disappear. Trials started with the low target, which remained on the screen until the end of the block unless hit by the cursor. Once the subject decreased the CN’s firing rate enough to drive the cursor into the low target, the high target immediately appeared. The subject then had a 2-s window to drive the cursor into the high target. If the subject accomplished this, the high target disappeared and a liquid reward was made available, and the low target immediately reappeared to begin the next trial. If the subject did not hit the high target within the 2-s window, the high target disappeared without a reward and the low target immediately reappeared, beginning the next trial. For each block, Monkeys N and R were given 4 min to complete as many trials as possible in the given sensory context, with the exception of the FAST focused training paradigm. Monkey A’s blocks ranged in duration (1.54 ± 0.57 min, mean ± SD).

The CN’s instantaneous firing rate was extracted from spike counts binned at 30 Hz and smoothed with a 20-bin boxcar filter to reduce cursor jitter. This filtered firing rate *r_f_* was translated into a 1D cursor position s along the movement axis according to the linear map

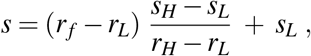

where *r_H_* and *r_L_* are the firing rates that the CN’s firing had to reach in order for the cursor to enter the high and low targets, respectively. The 1D positions of those targets along the movement axis, *s_H_* and *s_L_*, were always set to be 34 mm to either side of the axis origin.

In the FAST paradigm, sensory context is defined by the orientation of the movement axis and the location of the axis origin within the display. Sensory context changed only between blocks, such that each block of trials corresponded only to a single sensory condition.

#### FAST orientation paradigm

In the FAST orientation experiments, each condition was characterized by the angle of the movement axis, with the axis origin fixed at the center of the display. We tested 8 orientations in 45° intervals. To control for motivation and other potential time-related confounds, in a typical FAST orientation session each angle was presented twice, non-consecutively, for a total of 16 blocks of trials. Orientations were ordered randomly but constrained to ensure that all conditions were presented once before any were presented for the second time, and to prevent *mirrored* orientations (i.e., angles separated by exactly 180°, which produce parallel movement axes with opposite polarity) from appearing consecutively.

#### FAST location paradigm

In the FAST location experiments, we fixed axis orientation and instead varied the location of the axis origin in the workspace (Fig. 5a). To extend this beyond a single orientation, we presented each location at a second orientation as well. Conditions were therefore defined as a location-orientation combination. We trained two subjects in this paradigm. In addition to the center of the display (O), we defined eight other possible axis locations, each 34 mm from the center: four following the cardinal directions (North, N; West, W; South, S; East, E) and another four interspersed between them (Northwest, NW; Southwest, SW; Southeast, SE; Northeast, NE). For Monkey N, we tested each CN at five axis locations (O, NW, SW, SE, NE) across two orientations, resulting in 10 location-orientation conditions, and allowed two blocks per combined condition. For Monkey R, we tested each CN at all nine locations (O, N, NW, W, SW, W, SE, E, NE) across two orientations, resulting in 18 location-orientation conditions, but provided only one block per combination.

To establish a controllability baseline, Monkey N was given four blocks at the start of each session that combined the central location (O) with four angles: two arbitrary orientations and their mirrored counterparts. We selected two orientations—one from each mirrored pair—in which the subject had performed well and tested all five axis locations at the two selected orientations, such that each location-orientation condition would be presented twice in total. Blocks were ordered randomly but constrained to ensure that all location-orientation conditions were presented once before any were presented for the second time, and that both the orientation and the location of the movement axis changed across sequential blocks.

For Monkey R, each location-orientation condition appeared only once and in an order that ensured that location changed across sequential blocks but blocks were grouped by orientation: first all nine locations were randomly presented at one constant orientation, then all locations were randomly presented at the second orientation.

#### FAST focused training paradigm

The focused training paradigm tested controllability across orientations twice: once before and once after the *focus* blocks (Fig. 6a). Similarly to the FAST orientation paradigm, each focused training experiment started with a single 4-min block at each of the 8 orientations. This served to establish a baseline level of controllability across all conditions. We then selected one of the orientations (typically, an “average performance” condition, neither the best nor the worst) and ran one 15-min block at this orientation. As before, the difficulty ratcheting was on throughout this focus block, such that the firing rate targets would gradually separate if performance was strong. At the completion of this 15-min block, the subject was given a new 15-min focus block at that same orientation. This served to reset trial difficulty between focus blocks and make the task slightly easier again, renewing the monkey’s motivation. Once this second 15-min block was completed, we ran a second shuffled set of 4-min blocks across all 8 orientation conditions.

#### FAST multi-day training paradigm

We developed the multi-day training paradigm to test whether continued practice over several sessions would eliminate the interaction between sensory context and controllability. In a FAST multi-day training experiment, day 1 was identical to a typical FAST orientation session. We then held both the CN and the initial target rate values constant for days 2 through 5 of the experiment and had the subject perform an equivalent FAST orientation session each day using those fixed parameters. In other words, other than by the order in which orientations were presented—which was randomized at the beginning of each day—the task was identical across all 5 days of the experiment. The CN was tracked across days by visual inspection, and confirmed offline through a neuron-tracking algorithm^62^.

#### Initializing FAST firing rate target values

Before running any of the FAST paradigms each day, the subjects first performed a short, standard BCI center-out session to collect baseline firing statistics from the neural population, which was used to set the FAST parameters. In this session, the subjects used a linear decoder^63^ to control the 2D velocity of a cursor in order to hit one of eight targets uniformly spaced around a circle of radius 85 mm. At the start of each trial, cursor position was reset to the center and a peripheral target appeared on the workspace. In a successful trial, the subject would drive the cursor to the target within 3 s and hold it in place for 100–300 ms to receive a reward. After decoder calibration, subjects performed 162 ± 60 trials (mean ± SD) in the task.

At the end of this BCI center-out session, we randomly selected one well-sorted neuron to be used as CN in the FAST paradigm, with a bias towards selecting a new neuron each day and choosing neurons that displayed a wide dynamic range. The FAST initial firing rate targets *r_H_* and *r_L_* were set, respectively, to the 80th and 20th percentiles of the firing rates exhibited by the selected CN throughout the BCI center-out session. The FAST paradigm enforced at all times a minimum difference between firing rate targets of 15 Hz and a minimum *r_L_* value of 3 Hz (i.e., at no point was the subject required to drive the CN’s firing rate below 3 Hz).

#### Adaptive difficulty schedule

For Monkeys N and R, the FAST paradigm included an automated difficulty schedule similar to the setup used by Schieber and colleagues^12^. Within each block, this adaptive mechanism would trigger a difficulty increment when it detected that recent task performance exceeded a certain threshold. Task difficulty is determined by the separation between the firing rate targets *r_H_* and *r_L_*. If the monkey was highly successful, these target rate values would be pushed farther apart, increasing the dynamic range required to reach the visual targets on the screen (Fig. 2a). Firing rate targets were reset to their original values at the start of every block.

The scale of the difficulty increments was relative to the CN’s firing statistics. Let *D* be the average of the 80th and 20th firing rate percentiles. Every 10 s, we computed the number of successes accrued over the past 20 s. If the number of successes was greater than 10 (i.e., greater than 30 successes/min), a full-step increase in difficulty was triggered, which increased the separation of the target rates by 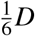. That is, *r_H_* was increased to 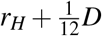 and *r_L_* was decreased to 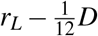. If the number of successes was not as high but was still greater than 7 (i.e., greater than 20 successes/min), a half-step increase in difficulty would instead be triggered, which increased the separation of the firing rate targets by 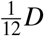. For the half-step increase, *r_H_* increased to 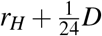, and *r_L_* decreased to 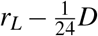. Note that if the change would decrease *r_L_* below 3 Hz, *r_L_* was instead left at its original value and the entirety of the change was instead applied to *r_H_*.

### Data Analysis

#### Normalization of the CN’s firing rate and its time derivative

Single-neuron firing rates *z_t_* were obtained offline by smoothing spiking event timestamps with a Gaussian kernel (*σ_k_* = 150 ms) and z-scoring these smoothed values over the entire FAST session for that day.

The normalized time derivative of the firing rate 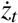 was computed by taking the finite difference of the smoothed, normalized firing rate over the sampling interval *h* =1 ms, and scaling by the Gaussian kernel width *σ_k_*:

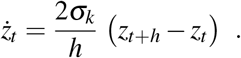

#### Computing controllability

We designed *controllability* as a metric that captures the ease with which the subject can voluntarily modulate the firing rate of the CN across its dynamic range, given a sensory feedback condition. In each block, we first identified the *peak minute*: the continuous minute of control that had the highest number of successful trials (Fig. 2b). We subdivided this peak minute into 12 non-overlapping, 5-s bins and computed each bin’s *control ellipse*: the 2-standard-deviation covariance ellipse between the CN’s normalized firing rates *z* and their time derivative 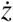. The area of this ellipse captures both the dynamic range achieved by the CN during this 5-s bin and the speed with which the subject was able to modulate it. We thus quantified the controllability of the CN, given a sensory context, as the average area of these control ellipses across the 12 bins of the peak minute (Fig. 2c-d). Blocks in which the subject failed to complete at least 5 successful trials within a continuous minute (or a minimum of 4 successful trials in the case of the FAST multi-day paradigm) were discarded during analysis. If a CN had more than two conditions not represented because of this performance threshold, it was removed due to insufficient data. Monkey A had 16 blocks shorter than 1 min (46 ± 8 s); instead of selecting a peak minute, in these cases the full duration of the block was binned at 5-s intervals and used to compute controllability values.

#### Statistical testing

We quantified the significance of the impact that sensory context had on a CN’s controllability using one-way ANOVA to partition the total variation found in controllability into across-condition variation and within-condition variation. The ratio between these components determines whether controllability means are significantly different across conditions. To assess the magnitude of this effect, we computed the fraction of variance present in controllability that could be explained by sensory context.

For each subject, we also tested whether the portion of variance explained by sensory context across neurons was statistically different from chance levels. To do this, we created a null distribution by shuffling the condition labels of every controllability value produced by each CN and computed the portion of variance in the CN’s shuffled data that was explained by sensory condition. Then, we used a two-sided Wilcoxon rank sum test to compare this *shuffled* distribution of explained variances with the subject’s original distribution of variances explained by sensory context across CNs. Comparisons in the FAST focused and multi-day training paradigms were calculated employing paired-sample t-tests.

#### Controlling for interactions between time and controllability

Since sensory conditions were run block-wise, we performed a control to test whether time might instead explain the relationship between controllability and sensory context in the FAST orientation, location and multi-day training paradigms. For each CN, we first fit a linear regression against block number to identify trends in controllability based on the chronological order in which the conditions had been given to the subject. We then removed that temporal effect by taking the residuals of the regression and computing the portion of variance in those residual controllability values that was explained by sensory context, following the same procedure described above for the computation and statistical testing of variance explained. The results of these analyses are presented in Supplementary Fig. 2, Supplementary Fig. 4 and Supplementary Fig. 7.

## DATA AVAILABILITY

The data that support the findings of this study are available from the corresponding author upon reasonable request.

## AUTHOR CONTRIBUTIONS

C.F.F. and S.M.C. conceived and designed research. C.F.F. performed experiments and analyzed data. C.F.F. and S.M.C. interpreted results of experiments. C.F.F. and S.M.C. drafted and revised manuscript. C.F.F and S.M.C. approved final version of manuscript.

## Notes

### Competing Interest Statement

The authors have declared no competing interest.

## REFERENCES

1. Porter, R. & Lemon, R. Corticospinal Function and Voluntary Movement (Oxford University Press, Sept. 1993).

2. Golub, M. D., Chase, S. M., Batista, A. P. & Yu, B. M. Brain–computer interfaces for dissecting cognitive processes underlying sensorimotor control. Current Opinion in Neurobiology 37 (2016).

3. Fetz, E. E. Operant Conditioning of Cortical Unit Activity. Science 163 (1969).

4. Fetz, E. E. & Baker, M. A. Operantly conditioned patterns on precentral unit activity and correlated responses in adjacent cells and contralateral muscles. Journal of Neurophysiology 36 (Mar. 1973).

5. Schieber, M. H. Dissociating motor cortex from the motor. The Journal of physiology 589 (2011).

6. Davidson, A. G., Chan, V., O’Dell, R. & Schieber, M. H. Rapid changes in throughput from single motor cortex neurons to muscle activity. Science (2007).

7. Fetz, E. E. & Finocchio, D. V. Correlations between activity of motor cortex cells and arm muscles during operantly conditioned response patterns. Experimental Brain Research 23 (Sept. 1975).

8. Taylor, D. M., Tillery, S. I. H. & Schwartz, A. B. Direct Cortical Control of 3D Neuroprosthetic Devices. Science 296 (June 2002).

9. Carmena, J. M. et al. Learning to Control a Brain–Machine Interface for Reaching and Grasping by Primates. PLoS Biology 1 (Oct. 2003).

10. Moritz, C. T., Perlmutter, S. I. & Fetz, E. E. Direct control of paralysed muscles by cortical neurons. Nature 456 (Dec. 2008).

11. Moritz, C. T. & Fetz, E. E. Volitional control of single cortical neurons in a brain-machine interface. Journal of neural engineering 8 (2011).

12. Law, A. J., Rivlis, G. & Schieber, M. H. Rapid acquisition of novel interface control by small ensembles of arbitrarily selected primary motor cortex neurons. Journal of Neurophysiology (2014).

13. Koralek, A. C., Jin, X., Long II, J. D., Costa, R. M. & Carmena, J. M. Corticostriatal plasticity is necessary for learning intentional neuroprosthetic skills. Nature 483 (2012).

14. Kennedy, P., Bakay, R., Moore, M., Adams, K. & Goldwaithe, J. Direct control of a computer from the human central nervous system. IEEE Transactions on Rehabilitation Engineering 8 (June 2000).

15. Rosén, I. & Asanuma, H. Peripheral afferent inputs to the forelimb area of the monkey motor cortex: Input-output relations. Experimental Brain Research 14 (1972).

16. Sato, T. R. & Svoboda, K. The Functional Properties of Barrel Cortex Neurons Projecting to the Primary Motor Cortex. Journal of Neuroscience 30 (Mar. 2010).

17. Hooks, B. M. et al. Organization of Cortical and Thalamic Input to Pyramidal Neurons in Mouse Motor Cortex. Journal of Neuroscience 33 (2013).

18. Kleinfeld, D., Sachdev, R. N., Merchant, L. M., Jarvis, M. R. & Ebner, F. F. Adaptive Filtering of Vibrissa Input in Motor Cortex of Rat. Neuron 34 (June 2002).

19. Ferezou, I. et al. Spatiotemporal Dynamics of Cortical Sensorimotor Integration in Behaving Mice. Neuron 56 (Dec. 2007).

20. Huber, D. et al. Multiple dynamic representations in the motor cortex during sensorimotor learning. Nature 484 (Apr. 2012).

21. Petrof, I., Viaene, A. N. & Sherman, S. M. Properties of the primary somatosensory cortex projection to the primary motor cortex in the mouse. Journal of neurophysiology 113 (2015).

22. Stavisky, S. D., Kao, J. C., Ryu, S. I. & Shenoy, K. V. Motor Cortical Visuomotor Feedback Activity Is Initially Isolated from Downstream Targets in Output-Null Neural State Space Dimensions. Neuron 95 (July 2017).

23. Suminski, A. J., Tkach, D. C., Fagg, A. H. & Hatsopoulos, N. G. Incorporating Feedback from Multiple Sensory Modalities Enhances Brain-Machine Interface Control. Journal of Neuroscience 30 (Dec. 2010).

24. Bedingham, W. & Tatton, W. Dependence of EMG Responses Evoked by Imposed Wrist Displacements on Preexisting Activity in the Stretched Muscles. Canadian Journal of Neurological Sciences /Journal Canadien des Sciences Neurologiques 11 (May 1984).

25. Pruszynski, J. A. & Scott, S. H. Optimal feedback control and the long-latency stretch response. Experimental Brain Research 218 (2012).

26. Day, B. L. & Lyon, I. N. Voluntary modification of automatic arm movements evoked by motion of a visual target. Experimental Brain Research 130 (Jan. 2000).

27. Franklin, D. W. & Wolpert, D. M. Specificity of Reflex Adaptation for Task-Relevant Variability. Journal of Neuroscience 28 (Dec. 2008).

28. Evarts, E. V. & Tanji, J. Reflex and intended responses in motor cortex pyramidal tract neurons of monkey. Journal of Neurophysiology 39 (Sept. 1976).

29. Cheney, P. D. & Fetz, E. E. Corticomotoneuronal cells contribute to long-latency stretch reflexes in the rhesus monkey. The Journal of Physiology 349 (Apr. 1984).

30. Jacobs, J. V. & Horak, F. B. Cortical control of postural responses. Journal of Neural Transmission 114 (Oct. 2007).

31. Scott, S. H. Optimal feedback control and the neural basis of volitional motor control. Nature Reviews Neuroscience 5 (July 2004).

32. Pruszynski, J. A. Primary motor cortex and fast feedback responses to mechanical perturbations: a primer on what we know now and some suggestions on what we should find out next. Frontiers in Integrative Neuroscience 8 (Sept. 2014).

33. Scott, S. H., Cluff, T., Lowrey, C. R. & Takei, T. Feedback control during voluntary motor actions. Current Opinion in Neurobiology 33 (Aug. 2015).

34. Downey, J. E. et al. Motor cortical activity changes during neuroprosthetic-controlled object interaction. Scientific Reports 7 (Dec. 2017).

35. Hennig, J. A. et al. Learning is shaped by abrupt changes in neural engagement. Nature Neuroscience 24 (May 2021).

36. Stavisky, S. D. et al. Brain-machine interface cursor position only weakly affects monkey and human motor cortical activity in the absence of arm movements. Scientific Reports 8 (Dec. 2018).

37. Scott, S. H. & Kalaska, J. F. Reaching Movements With Similar Hand Paths But Different Arm Orientations. I. Activity of Individual Cells in Motor Cortex. Journal of Neurophysiology 77 (Feb. 1997).

38. Sergio, L. E. & Kalaska, J. F. Systematic Changes in Motor Cortex Cell Activity With Arm Posture During Directional Isometric Force Generation. Journal of Neurophysiology 89 (Jan. 2003).

39. Wilson, S. M., Saygin, A. P., Sereno, M. I. & Iacoboni, M. Listening to speech activates motor areas involved in speech production. Nature Neuroscience 7 (July 2004).

40. Cheung, C., Hamilton, L. S., Johnson, K. & Chang, E. F. The auditory representation of speech sounds in human motor cortex. eLife 5 (Mar. 2016).

41. Rouse, A. G. & Schieber, M. H. Spatiotemporal Distribution of Location and Object Effects in Primary Motor Cortex Neurons during Reach-to-Grasp. Journal of Neuroscience 36 (Oct. 2016).

42. Ruff, D. A. & Cohen, M. R. Simultaneous multi-area recordings suggest that attention improves performance by reshaping stimulus representations. Nature Neuroscience 22 (Oct. 2019).

43. Steinmetz, N. A., Zatka-Haas, P., Carandini, M. & Harris, K. D. Distributed coding of choice, action and engagement across the mouse brain. Nature 576 (Dec. 2019).

44. Smoulder, A. L. et al. Monkeys exhibit a paradoxical decrease in performance in high-stakes scenarios. Proceedings of the National Academy of Sciences 118 (Aug. 2021).

45. Perge, J. A. et al. Intra-day signal instabilities affect decoding performance in an intracortical neural interface system. Journal of Neural Engineering 10 (June 2013).

46. Dunlap, C. F., Colachis, S. C., Meyers, E. C., Bockbrader, M. A. & Friedenberg, D. A. Classifying Intracortical Brain-Machine Interface Signal Disruptions Based on System Performance and Applicable Compensatory Strategies: A Review. Frontiers in Neurorobotics 14 (Oct. 2020).

47. Cowley, B. R. et al. Slow Drift of Neural Activity as a Signature of Impulsivity in Macaque Visual and Prefrontal Cortex. Neuron 108 (Nov. 2020).

48. Sadtler, P. T. et al. Neural constraints on learning. Nature 512 (2014).

49. Golub, M. D. et al. Learning by neural reassociation. Nature Neuroscience 21 (Apr. 2018).

50. Oby, E. R. et al. New neural activity patterns emerge with long-term learning. Proceedings of the National Academy of Sciences 116 (July 2019).

51. Zhou, X., Tien, R. N., Ravikumar, S. & Chase, S. M. Distinct types of neural reorganization during long-term learning. Journal of Neurophysiology 121 (Apr. 2019).

52. Krakauer, J. W., Pine, Z. M., Ghilardi, M.-F. & Ghez, C. Learning of Visuomotor Transformations for Vectorial Planning of Reaching Trajectories. The Journal of Neuroscience 20 (Dec. 2000).

53. Cerf, M. et al. On-line, voluntary control of human temporal lobe neurons. Nature 467 (Oct. 2010).

54. Clancy, K. B., Koralek, A. C., Costa, R. M., Feldman, D. E. & Carmena, J. M. Volitional modulation of optically recorded calcium signals during neuroprosthetic learning. Nat Neurosci 17 (June 2014).

55. Neely, R. M., Koralek, A. C., Athalye, V. R., Costa, R. M. & Carmena, J. M. Volitional Modulation of Primary Visual Cortex Activity Requires the Basal Ganglia. Neuron 97 (Mar. 2018).

56. Andersen, R. A., Aflalo, T. & Kellis, S. From thought to action: The brain–machine interface in posterior parietal cortex. Proceedings of the National Academy of Sciences 116 (Dec. 2019).

57. Patel, K., Katz, C. N., Kalia, S. K., Popovic, M. R. & Valiante, T. A. Volitional control of individual neurons in the human brain. Brain 144 (Dec. 2021).

58. Gallego, J. A., Makin, T. R. & McDougle, S. D. Going beyond primary motor cortex to improve brain–computer interfaces. Trends in Neurosciences 45 (Jan. 2022).

59. Fukuma, R. et al. Voluntary control of semantic neural representations by imagery with conflicting visual stimulation. Communications Biology 5 (Mar. 2022).

60. Jeon, B. B., Fuchs, T., Chase, S. M. & Kuhlman, S. J. Existing function in primary visual cortex is not perturbed by new skill acquisition of a non-matched sensory task. Nature Communications 13 (June 2022).

61. Velliste, M., Perel, S., Spalding, M. C., Whitford, a. S. & Schwartz, a. B. Cortical control of a robotic arm for self-feeding. Nature 453 (2008).

62. Fraser, G. W. & Schwartz, A. B. Recording from the same neurons chronically in motor cortex. Journal of Neurophysiology 107 (Apr. 2012).

63. Chase, S. M., Schwartz, A. B. & Kass, R. E. Bias, optimal linear estimation, and the differences between open-loop simulation and closed-loop performance of spiking-based brain–computer interface algorithms. Neural Networks 22 (Nov. 2009).

