## Supplementary Information for "Sensory constraints on volitional modulation of the motor cortex"

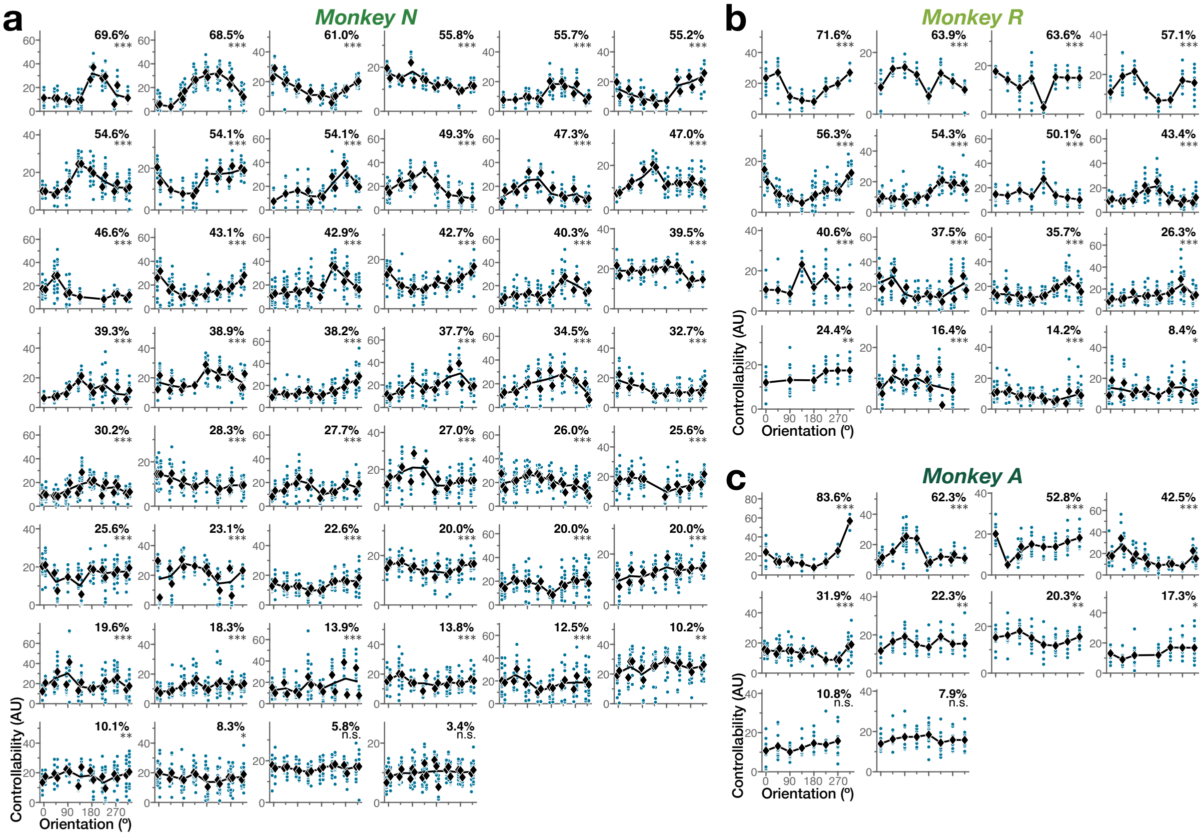


**Supplementary Fig. 1: All three subjects showed widespread orientation effects on their ability to volitionally modulate individual neurons in M1. a–c** Controllability across orientations for all CNs tested in the FAST orientation paradigm from Monkey N (**a**, $n = 46$ neurons), Monkey R (**b**, $n = 16$ neurons), and Monkey A (**c**, $n = 10$ neurons). For each neuron, individual controllability values (teal) and block averages (black) are shown; lines connect average controllability at each condition. Neurons are sorted by descending % variance explained by orientation, reported for each CN (ANOVA; $^{***}p < 0.001$, $^{**}p < 0.01$, $^{*}p < 0.05$, *n.s.* if $p \geq 0.05$).

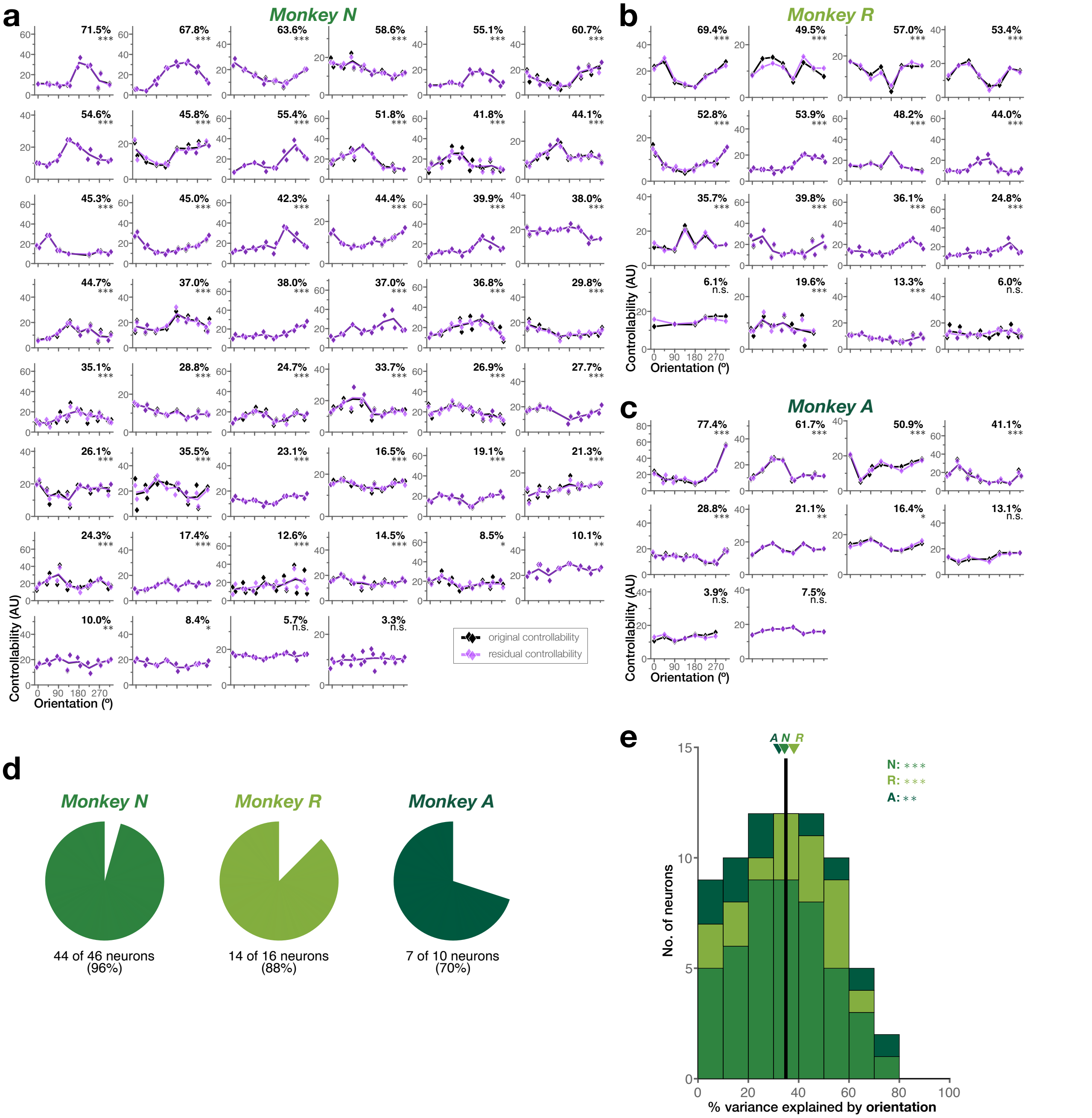


**Supplementary Fig. 2: The interaction between orientation and controllability cannot be explained as a linear effect of time. a–c** To assess linear effects of time, we fit for each CN a linear regression of controllability against block number. Then, we computed the residual controllability values from that regression to remove the linear effect caused by the passage of time throughout the session (see Methods for further details). Here we show how residual controllability (purple) compared to the original controllability values (black) across orientations for all CNs tested in the FAST orientation paradigm from Monkey N (**a**, $n = 46$ neurons), Monkey R (**b**, $n = 16$ neurons), and Monkey A (**c**, $n = 10$ neurons). Diamonds indicate block averages; lines connect average controllability at each condition. The portion of variance in residual controllability that is explained by orientation (%) is reported (ANOVA; $^{***}p < 0.001$, $^{**}p < 0.01$, $^{*}p < 0.05$, *n.s.* if $p \geq 0.05$). Neurons are sorted to correspond to Supplementary Fig. 1. **d** Fraction of CNs that display significant controllability differences given orientation after accounting for the passage of time (ANOVA, $p < 0.05$).**e** Distribution of the portion of variance in controllability explained by orientation after removing the effect of time. Vertical line indicates average across subjects; arrows indicate subject averages. The portion of variance explained by orientation across CNs continued to be statistically above chance for all subjects after controlling for the passage of time throughout a session (Wilcoxon rank sum test; N: $T = 3,175$, $p = 6.2 \times 10^{-16}$; R: $T = 380$, $p = 1.3 \times 10^{-5}$; A: $T = 144$, $p = 0.004$; $^{***}p < 0.001$, $^{**}p < 0.01$).

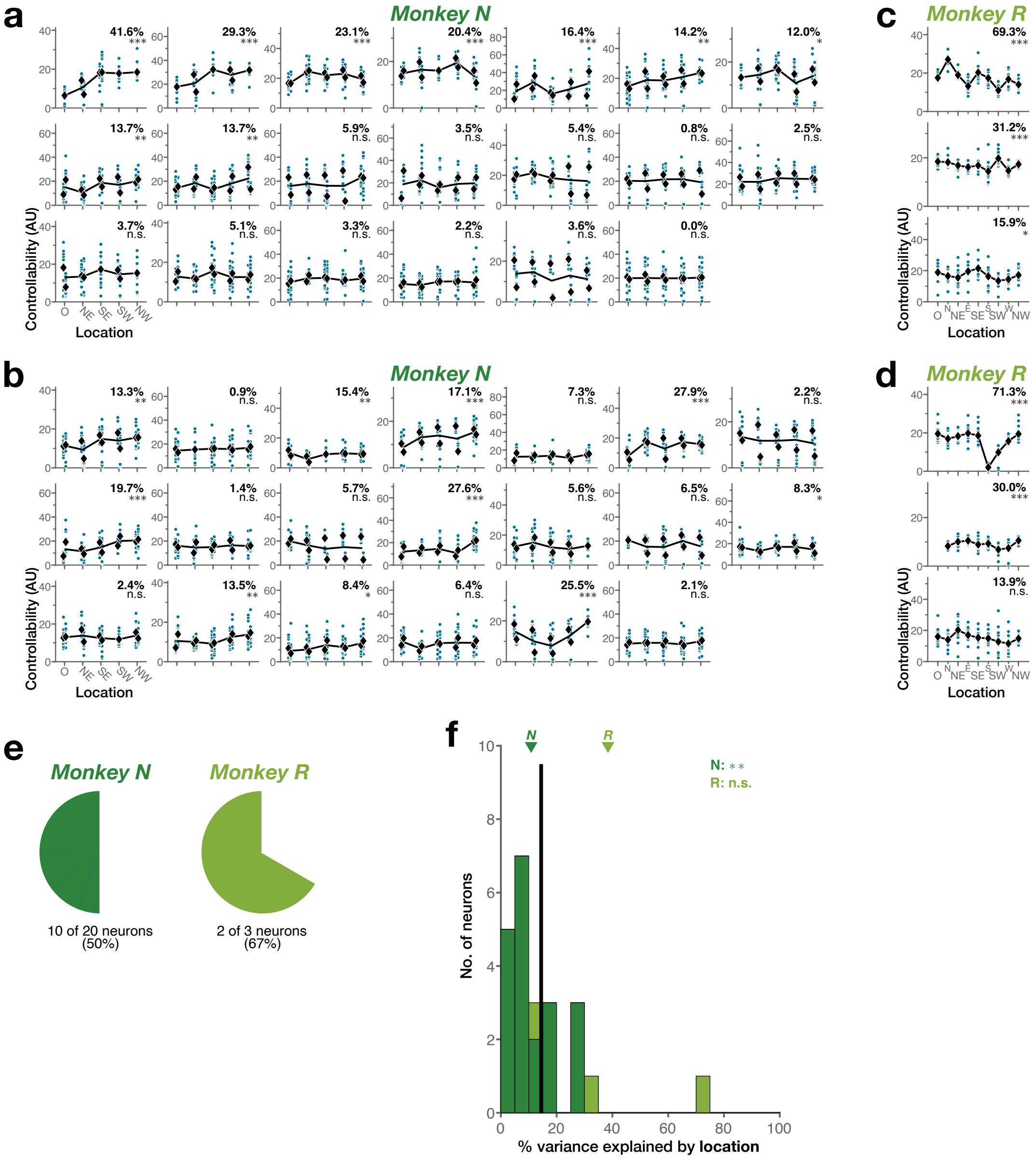


**Supplementary Fig. 3: The effects of location on controllability were comparable across the two orientations tested, and moderate compared to the impact of orientation. a–d** Controllability across all locations for Monkey N ($n$ = 20 neurons) at the first (**a**) and second (**b**) orientation, and for Monkey R ($n$ = 3 neurons) at the first (**c**) and second (**d**) orientation. Individual controllability values (teal) and block averages (black) are shown for each neuron; lines connect average controllability at each condition. Neurons are sorted by descending % variance explained by location at the first orientation, reported for each CN (one-way ANOVA; $***p < 0.001$, $*p < 0.05$, $n.s.\ p \geq 0.05$). **e** Fraction of CNs that display significant controllability differences given location on the second orientation (ANOVA, $p < 0.05$). **f** Distribution of the portion of variance in controllability explained by location, for the second orientation. Vertical line indicates average across subjects; arrows indicate subject averages. The portion of variance explained by location across CNs was statistically above chance for Monkey N but not Monkey R for the second orientation too (Wilcoxon rank sum test; N: $T = 513$, $p = 0.006$; R: $T = 15$, $p = 0.1$; $**p < 0.01$, $n.s.\ p \geq 0.05$).

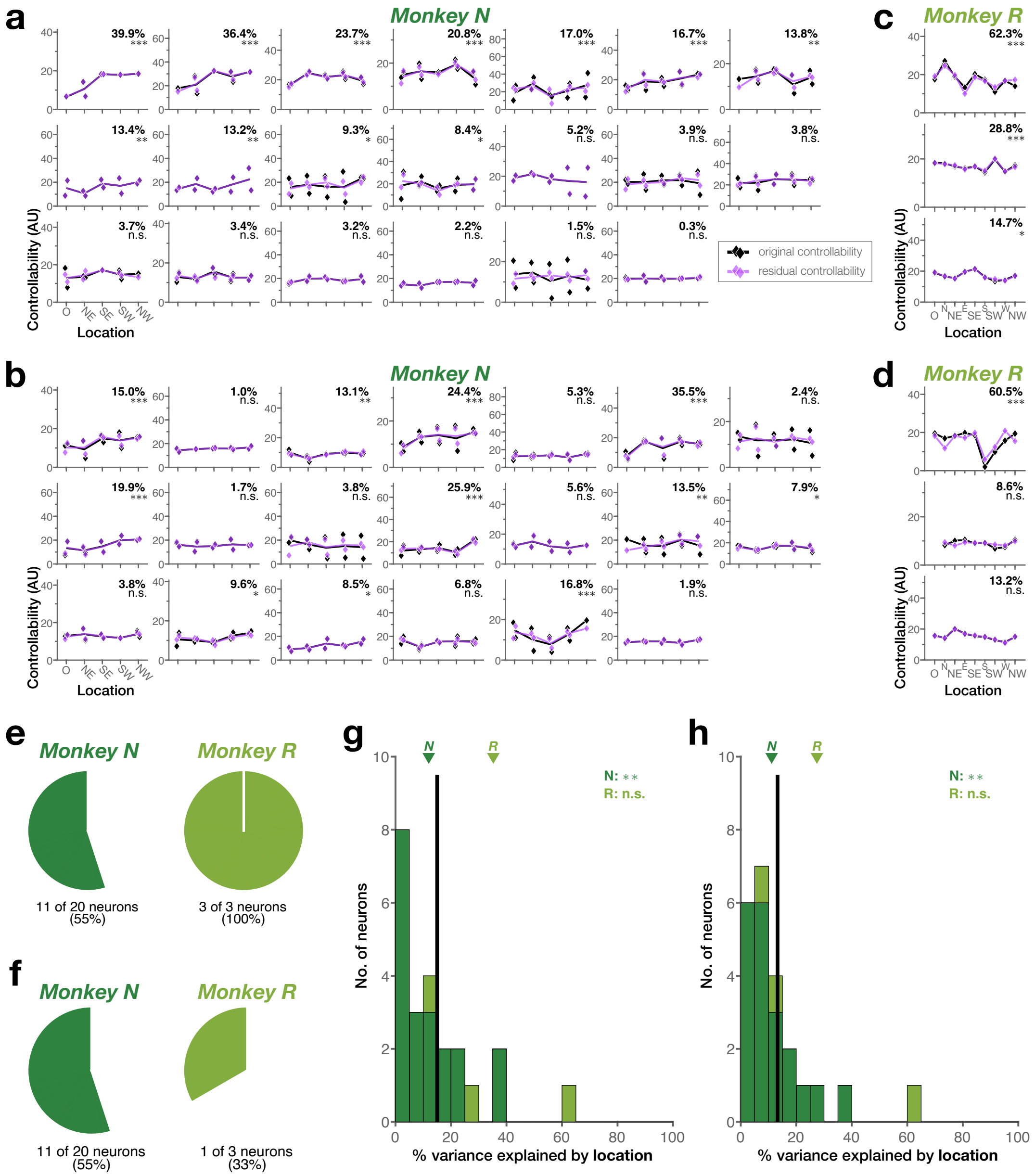

**Supplementary Fig. 4: The interaction between location and controllability cannot be explained as a linear effect of time for either orientation. a–d** Time-residual controllability across locations for Monkey N ($n = 20$ neurons) at the first (**a**) and second (**b**) orientation, and for Monkey R ($n = 3$ neurons) at the first (**c**) and second (**d**) orientation. Diamonds indicate block averages; lines connect average controllability at each condition. Residual controllability after removing the effects of time (purple) is shown together with controllability values (black), for comparison. The portion of variance in residual controllability that is explained by location (%) is reported (ANOVA; $***p < 0.001$, $**p < 0.01$, $*p < 0.05$, *n.s.* if $p \geq 0.05$). Neurons are sorted to correspond to Supplementary Fig. 3. **e, f** Fraction of CNs that display significant controllability differences given location after accounting for the passage of time, for the first (**e**) and second (**f**) orientation (ANOVA, $p < 0.05$). **g, h** Distribution of the portion of variance in controllability explained by location after removing the effect of time, for the first (**g**) and second (**h**) orientation. Vertical line indicates average across subjects; arrows indicate subject averages. For both orientations, the portion of variance explained by location across CNs continued to be statistically above chance for Monkey N but not Monkey R after controlling for the passage of time (Wilcoxon rank sum test; N: $T_{1st} = 515$, $p_{1st} = 0.005$, $T_{2nd} = 520$, $p_{2nd} = 0.003$; R: $T_{1st} = 15$, $p_{1st} = 0.1$, $T_{2nd} = 13$, $p_{2nd} = 0.4$; $**p < 0.01$, *n.s.* $p \geq 0.05$).

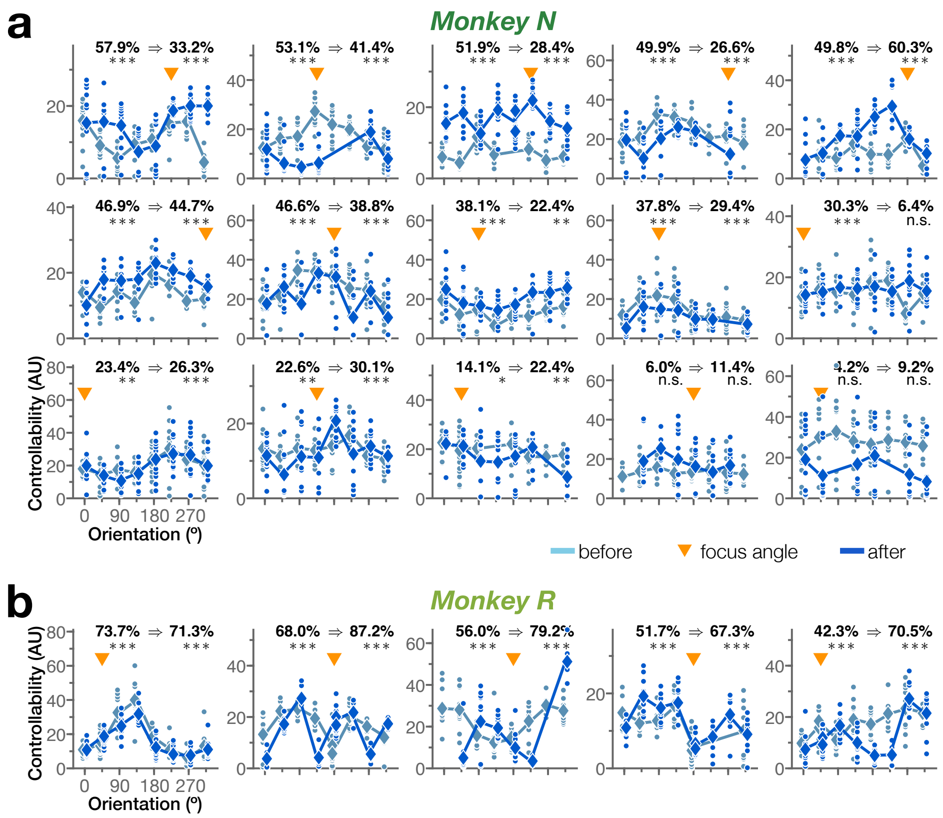


**Supplementary Fig. 5: Across neurons, focused training on challenging sensory contexts did not extinguish orientation dependence.** **a, b** Controllability across orientations before and after additional practice for all focused training CNs from Monkey N (**a**, $n = 15$ neurons) and Monkey R (**a**, $n = 5$ neurons). Controllability values (dots), block averages (diamonds) and lines connecting average controllability at each condition, before and after focus blocks, are shown for each CN. Variance explained by orientation is given as *before* ⇒ *after* (one-way ANOVA; $^{***}p < 0.001$, $^{**}p < 0.01$); neurons are sorted by descending variance explained before focused training.

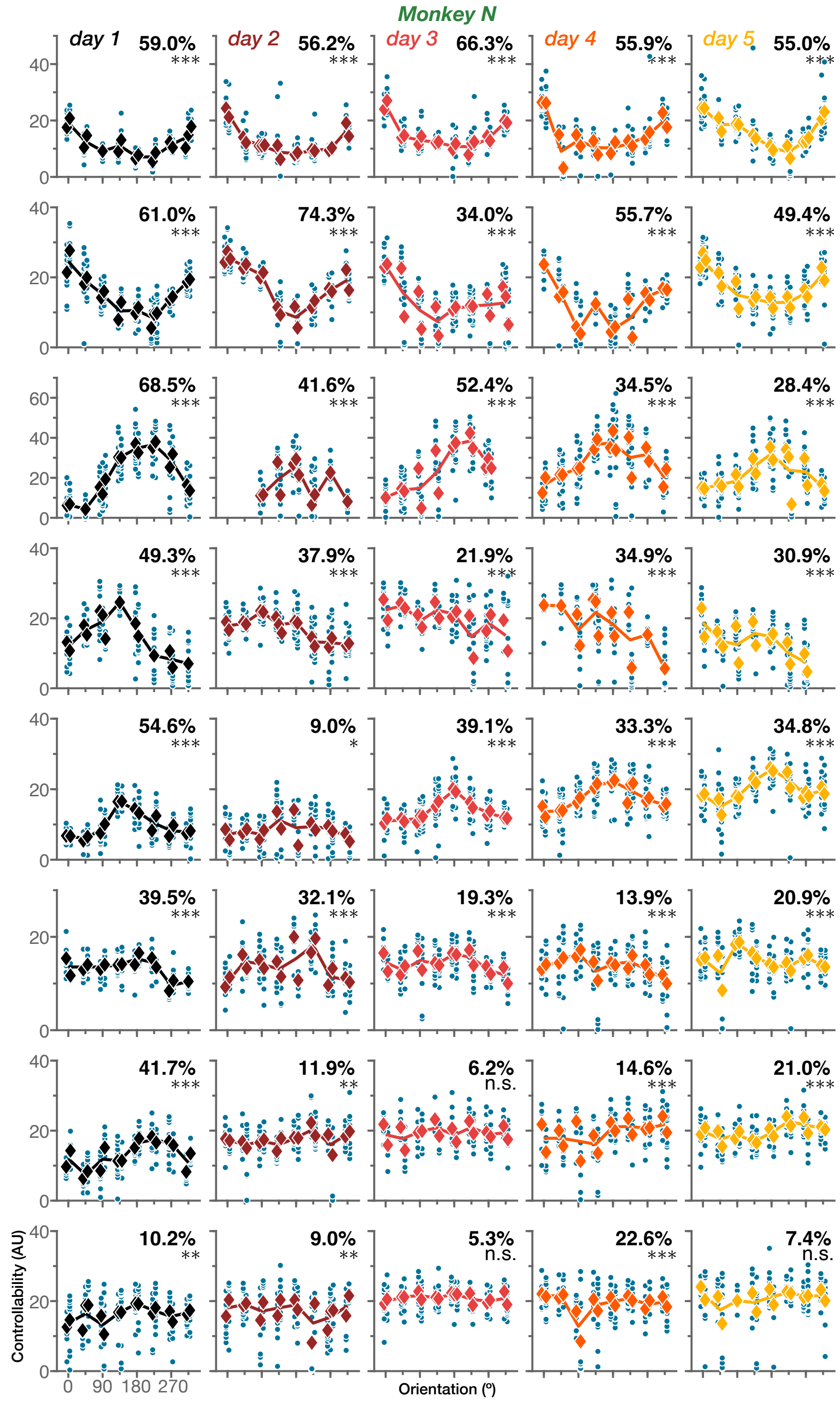


**Supplementary Fig. 6: Training over several days is not sufficient to extinguish orientation dependence.** Controllability across orientations throughout multiple days of training for all multi-day CNs from Monkey N ($n$ = 8 neurons). Individual controllability values (teal) are shown for each neuron (rows) and training day (columns); average controllability per block (diamond) and lines connecting average controllability at each condition are color-coded by training day. Neurons are sorted by descending average % variance explained by orientation across days, reported for each CN and day (one-way ANOVA; $***p < 0.001$, $**p < 0.01$, $*p < 0.05$, *n.s.* $p \geq 0.05$).

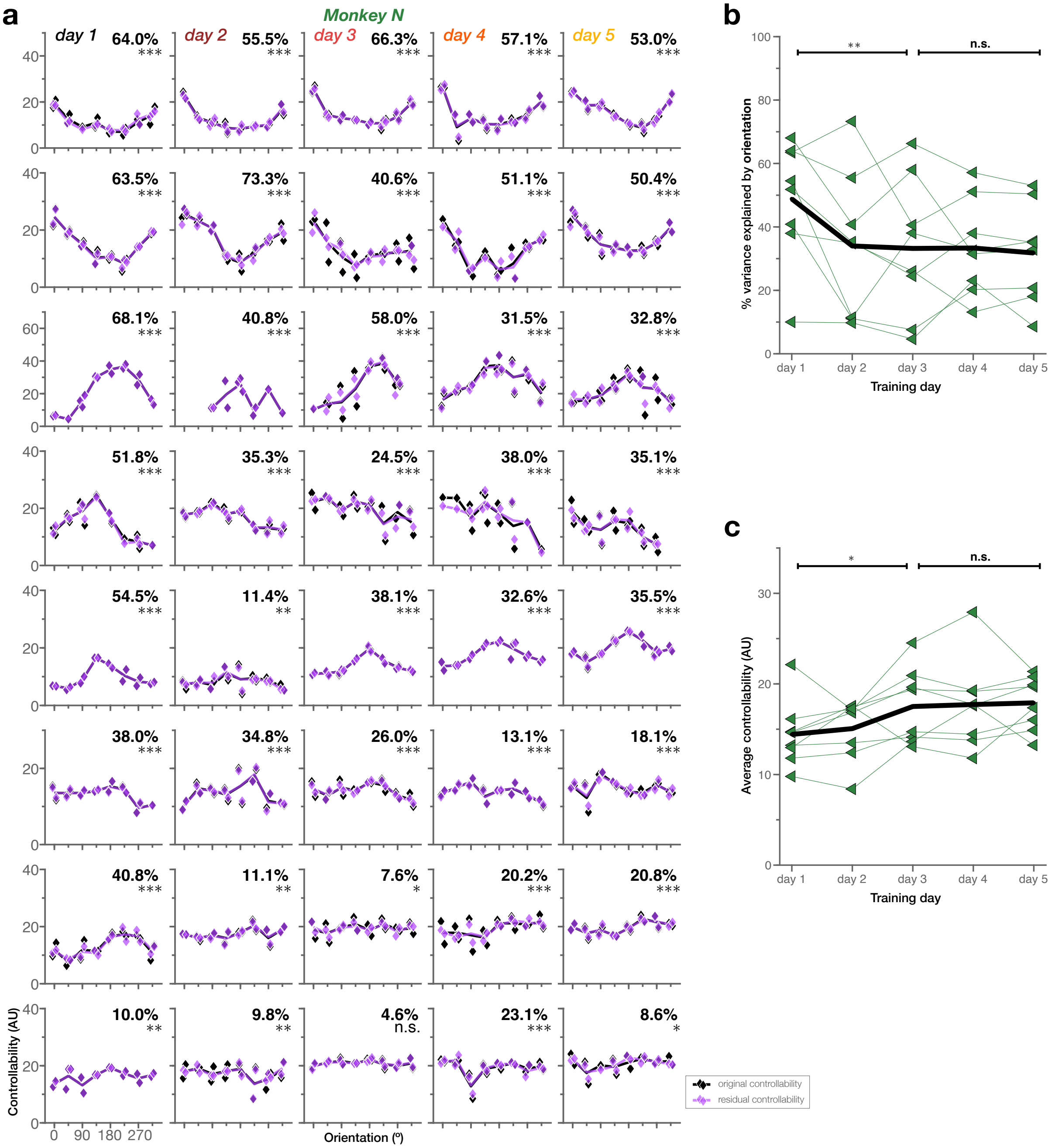


**Supplementary Fig. 7: The interaction between orientation and controllability across multiple days cannot be explained as a linear effect of time within each session. a** Day-wise time-residual controllability across orientations throughout multiple days of training for Monkey N ($n = 8$ neurons). Diamonds indicate block averages; lines connect average controllability at each condition. Residual controllability after removing the effects of time (purple) is shown together with original controllability values (black), for comparison. The portion of variance in residual controllability that is explained by orientation (%) is reported for each CN and day (ANOVA; $***p < 0.001$, $**p < 0.01$, $*p < 0.05$, *n.s.* if $p \geq 0.05$). Neurons are sorted to correspond to Supplementary Fig. 6 **b** Portion of variance in controllability that is explained by orientation across days, after removing the effect of time within each session. Lines represent the different CNs; black trace indicates average across CNs. Early training (days 1–3) and late training (days 3–5) comparisons are reported (paired-sample t-test; $**p < 0.01$, $*p < 0.05$, *n.s.* $p \geq 0.05$). **c** Average controllability across days, after removing the effect of time within each session. Same conventions as **(b)**. Overall, early training (days 1–3) continued to show significant differences in both % variance explained and average controllability after accounting for the passage of time within sessions, while late training (days 3–5) again did not (paired-sample t-test; $**p < 0.01$, $*p < 0.05$, *n.s.* $p \geq 0.05$).
